# Multiplexed Single-Molecule Epigenetic Analysis of Plasma-Isolated Nucleosomes for Cancer Diagnostics

**DOI:** 10.1101/2021.11.01.466724

**Authors:** Vadim Fedyuk, Nir Erez, Noa Furth, Olga Beresh, Ekaterina Andreishcheva, Abhijeet Shinde, Daniel Jones, Barak Bar Zakai, Yael Mavor, Tamar Peretz, Ayala Hubert, Jonathan E Cohen, Azzam Salah, Mark Temper, Albert Grinshpun, Myriam Maoz, Aviad Zick, Guy Ron, Efrat Shema

**Author notes:** Equal contribution (Vadim Fedyuk, Nir Erez).

## Abstract

The analysis of cell-free DNA (cfDNA) in plasma represents a rapidly advancing field in medicine, providing information on pathological processes in the body. Blood cfDNA is in the form of nucleosomes, which maintain their tissue- and cancer-specific epigenetic state. We developed EPINUC, a single-molecule multi-parametric assay to comprehensively profile the <u>E</u>pigenetics of <u>P</u>lasma <u>I</u>solated <u>Nuc</u>leosomes, DNA methylation and cancer-specific protein biomarkers. Our system allows high-resolution detection of six active and repressive histone modifications, their ratios and combinatorial patterns, on millions of individual nucleosomes by single-molecule imaging. In addition, it provides sensitive and quantitative data on plasma proteins, including detection of non-secreted tumor-specific proteins such as mutant p53. Applying this analysis to a cohort of plasma samples detected colorectal cancer at high accuracy and sensitivity, even at early stages. Finally, combining EPINUC with direct single-molecule DNA sequencing revealed the tissue-of-origin of colorectal, pancreatic, lung and breast tumors. EPINUC provides multi-layered clinical-relevant information from limited liquid biopsy material, establishing a novel approach for cancer diagnostics.

---

Non-invasive liquid biopsy methods, based on the analysis of cfDNA, potentiate a new generation of diagnostic approaches. The cfDNA that circulates in the plasma and serum of healthy individuals originates predominantly from death of normal blood cells^1^. In cancer patients, however, a fraction of cfDNA is tumor-derived, termed circulating tumor DNA (ctDNA). ctDNA-based sequence analysis has been shown to reveal tumor-specific genetic alterations and provide the means for non-invasive molecular profiling of tumors^2, 3^. Despite encouraging data, these approaches are limited, as they require genetic differences (i.e. mutations) in order to distinguish between the normal and tumor DNA. Liquid biopsy approaches based on analysis of non-genetic features have emerged recently, most prominently methodologies that utilize tissue- and cancer-specific DNA methylation, as well as differential fragmentation patterns of cfDNA^4–8^.

cfDNA in the plasma appears predominantly in the form of nucleosomes (cfNucleosomes), the basic unit of chromatin that consists of ∼150 base pairs of DNA wrapped around the octamer of core histone proteins. Histones are extensively modified by covalent attachment of various chemical groups, forming combinatorial epigenetic patterns that are unique to each tissue, and provide information on gene expression and regulatory elements within cells^9–12^. There is evidence that cfNucleosomes retain at least some of their epigenetic modifications, and a recent study applied Chromatin Immunoprecipitation and sequencing (ChIP-seq) to identify certain marks^13–15^. Moreover, deep sequencing of cfDNA revealed nucleosome occupancy patterns correlating with the tissue of origin^16–18^. While these approaches provide the first glimpse into the rich epigenetic information present in plasma that has so far remained mostly inaccessible, they have major limitations. Mainly, they require large amounts of input material, have a limited dynamic range (ChIP-seq), or are costly and require deep sequencing. Most importantly, these methodologies have limited output and sensitivity, as they usually measure a single layer of information (i.e. DNA methylation OR a single histone modification OR nucleosome occupancy, etc.). Thus, high-resolution approaches that integrate information from multiple parameters spanning different types of analytes are required.

Colorectal Cancer (CRC) is the third most common cancer worldwide, causing approximately 700,000 deaths every year^19^. Early metastatic seeding has been recently demonstrated in CRC^20^, underlining the necessity to develop better diagnostic tools to improve patient outcome. In this study, we developed a single-molecule-based liquid biopsy approach, to analyze multiple parameters from less than 1 ml of plasma sample and demonstrated its value for CRC diagnosis. We coined the technology “EPINUC” for Epigenetics of Plasma Isolated Nucleosomes (Fig. 1a). The technology builds on our recent development of a single-molecule system to image combinatorial histone modifications by Total Internal Reflection (TIRF) microscopy^21^. TIRF provides a powerful means for detecting single fluorescent molecules that are within ∼100 nm of a solid surface; a light source creates an evanescent wave that decays exponentially in intensity, thus eliminating background fluorescence from outside the focal plane. To capture nucleosomes from plasma, we developed high-efficiency enzymatic reactions to fluorescently tag and polyadenylate nucleosomes (Supplementary Fig.1a-e, Methods). Tailed, intact nucleosomes were then immobilized on a PEGylated surface via hybridization, and the status of their post-translational modifications (PTMs) was recorded by TIRF imaging with fluorescently tagged antibodies, verifying minimal spectral overlap (Fig. 1b and Supplementary Fig. 1f,g). Binding and dissociation of antibodies to target PTMs was imaged over 90 minutes, leveraging the TIRF narrow excitation range. Integration of binding events assured maximal detection of modified histones (Fig. 1c and Supplementary Fig. 2a).

**Fig.1:**
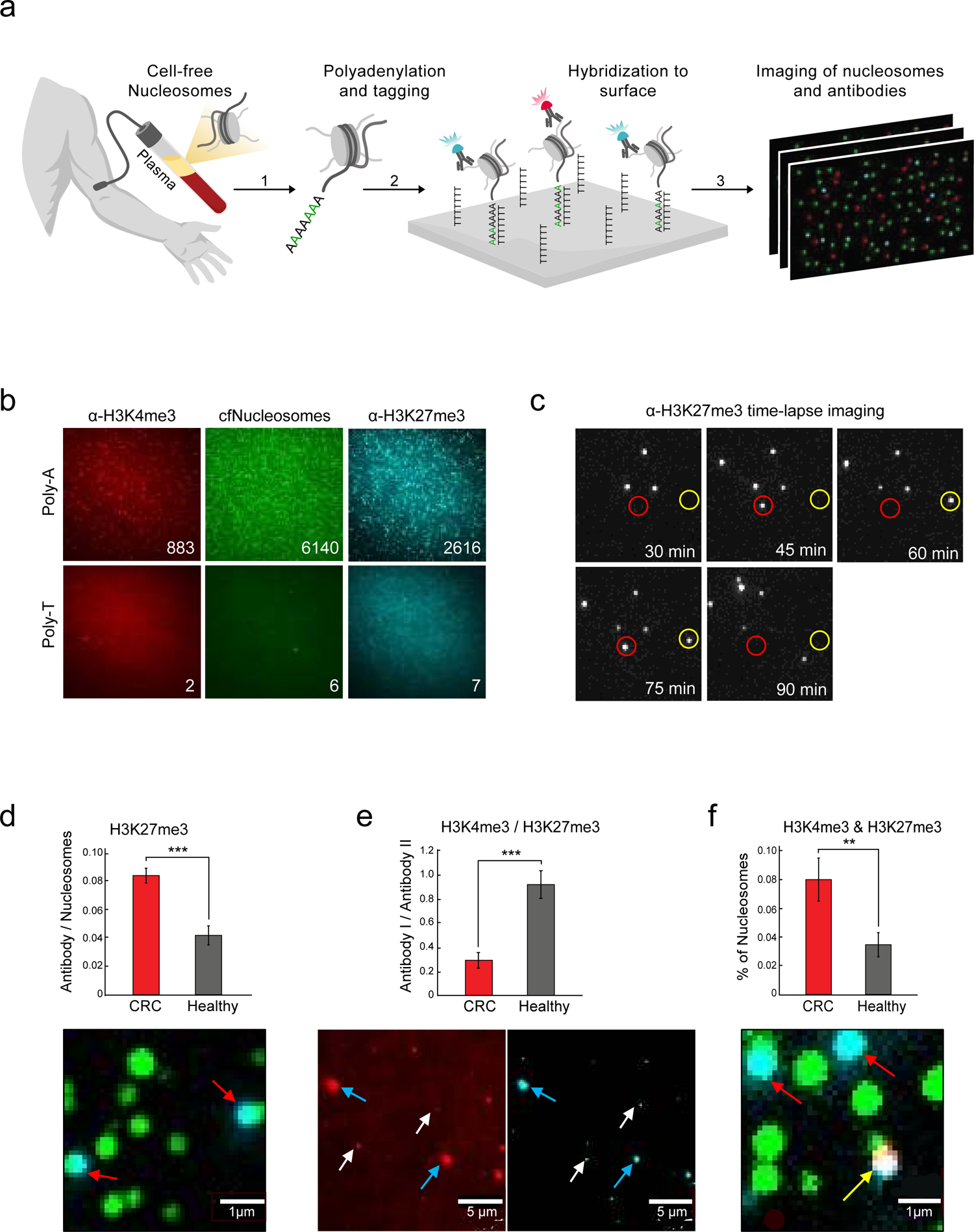
EPINUC decodes the combinatorial epigenetic states of plasma cell-free nucleosomes. **a,** Experimental scheme: (1) Sample preparation procedure of cfNucleosomes is carried out in one-step and consists of two enzymatic processes: repair of DNA ends by Klenow polymerase, and addition of a poly A tail by Terminal Transferase (TdT). The reaction contains a mixture of natural dATPs and fluorescently labeled dATPs (Cy3-dATP) to label nucleosomes. (2) cfNucleosmes are captured on a PEGylated-poly T surface via dA:dT hybridization. Immobilized nucleosomes are incubated with fluorescently labeled antibodies targeting different histone modifications. (3) TIRF microscopy is applied to record nucleosome positions and generate time-lapse imaging of antibodies’ binding events. **b,** Representative images of plasma-derived cfNucleosomes and the corresponding H3K4me3 and H3K27me3 signal (numbers within images represent counted spots). Each spot corresponds to a single nucleosome. Nucleosomes anchor to the surface specifically via hybridization, as evident from the lack of signal when tailed with dTTP (Poly-dT) rather than dATP (Poly-dA). **c,** Representative images of antibodies’ binding and dissociation events over time from individual target molecules (marked by red/yellow circles). **d,e,f** Example for quantification and representative images of the various parameters measured by EPINUC in plasma samples from one healthy subject and one CRC patient. Zoomed-in image segments of entire field of view (148µm^2^). **d,** The percentage of cfNucleosomes (green spots, Cy3) that are modified by H3K27me3 (cyan spots, AF488). Red arrows indicate co-localization events. Scale bar = 1µm **e,** Ratio between H3K4me3 (red, AF647) and H3K27me3 (cyan) antibodies. White arrows indicate antibody spots, blue arrows indicate TetraSpeck beads that are used for alignment. Scale bar = 5µm. **f,** cfNucleosomes marked by the combinatorial pattern of both H3K27me3 and H3K4me3. Red arrows indicate co-localization events of H3K27me3 only, yellow arrow indicates a combinatorially modified nucleosome. Scale bar = 1µm.

EPINUC relies on direct counting of single-molecules in a population, yielding data amenable to absolute quantification and comparisons between samples. Each antibody was verified for specificity and linearity of binding with a panel of recombinant modified nucleosomes, yielding six antibodies that passed the quality control criteria (Supplementary Fig. 2b-d). These antibodies target the tri-methylations on histone H3 lysine 9 (H3K9me3) and lysine 27 (H3K27me3), associated with gene silencing and heterochromatin, as well as antibodies targeting marks associated with active transcription: tri-methylation of histone H3 on lysine 4 (H3K4me3) and lysine 36 (H3K36me3), and acetylation on lysine 9 (H3K9ac). In addition, our panel includes an antibody targeting mono-methylation of histone H3 on lysine 4 (H3K4me1), a mark associated with enhancers^22, 23^.

Nucleosomes from each plasma sample were tagged with Cy3 (green), and imaged with three pairwise combinations of antibodies labeled with AF488 (cyan) or AF647 (red). Thus, we obtained multi-parametric data for six histone PTMs, comprising of the percentage of modified nucleosomes in each sample, the ratio between various histone modifications, and the percentage of nucleosomes that harbor a combinatorial pattern of two modifications (Fig. 1d-f). Repeated measurements of the same plasma samples produced highly similar results, attesting to the quantitative nature of this technology (Supplementary Fig.3). To the best of our knowledge, EPINUC is the only technology that enables counting of multiple histone PTMs, as well as combinatorially-modified nucleosomes, at a single-molecule precision, from low volume plasma sample (<1ml).

To extend the number of analytes measured beyond histone PTMs, we exploited the single-molecule system for quantification of protein biomarkers. We modulated surface chemistry to contain PEG-streptavidin, allowing anchoring of biotin-conjugated antibodies that target plasma proteins. Following incubation with plasma, bound proteins are imaged by fluorescent detection antibodies. Multiplexed simultaneous detection of three biomarkers is achieved through the use of distinct fluorophores (Fig. 2a). We imaged two proteins known to increase in plasma of CRC patients: Carcinoembryonic antigen (CEA), a canonical biomarker measured routinely by clinicians^24^, and Tissue inhibitor of metalloproteinase-1 (TIMP-1), a glycoprotein reported to have diagnostic value in screening for CRC^25^. In addition, we measured the mammalian sterile 20-like kinase 1 (MST1), an inhibitor of cell proliferation that decreases in CRC patients^26^ (Fig. 2b). We verified linear detection and specificity using cell-culture systems and knockdown experiments (Fig. 2c,d and Supplementary Fig. 4).

**Fig.2:**
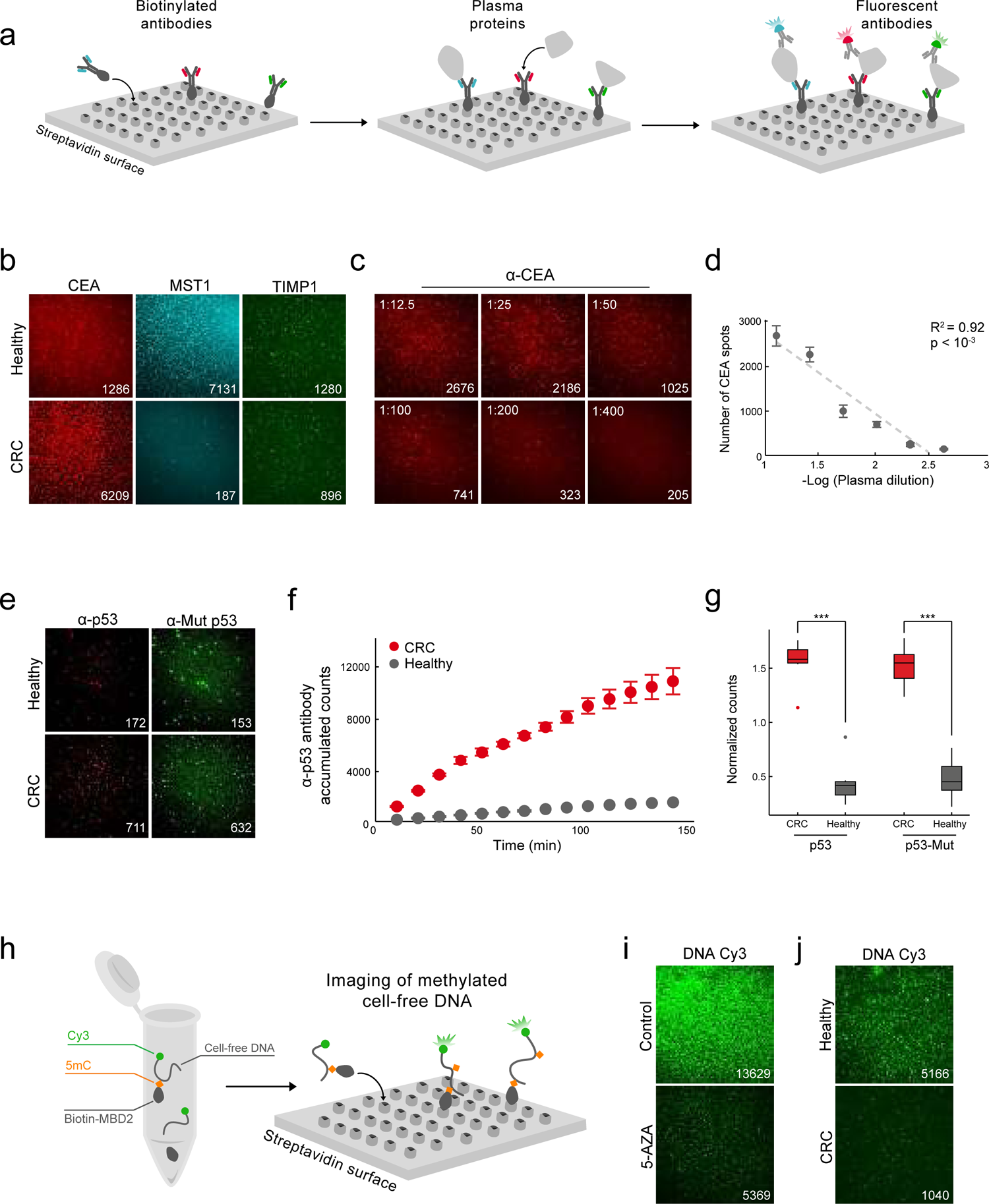
Multiplexed single-molecule detection of cancer-associated protein biomarkers, mutant p53 and DNA methylation. **a,** Experimental scheme: biotinylated capture antibodies targeting distinct proteins are anchored to a PEG-streptavidin surface. Plasma proteins are captured on surface, followed by detection with fluorescently labeled antibodies and TIRF imaging. Multiplexed detection of up to three proteins is achieved by labeling antibodies with different fluorophores. Each spot represents a single protein bound on the surface. **b,** Representative TIRF images of selected CRC biomarkers measured simultaneously for each plasma sample: CEA (red), MST1 (cyan) and TIMP-1(green). Images reveal distinct biomarkers profiles for healthy and CRC. **c,** Representative TIRF images depicting α-CEA antibody binding events (spots) over serial plasma dilutions. **d,** Regression analysis of the number of spots as a function of plasma concentration highlights the linearity of detection. Data is presented as the mean +/- s.d. of 50 FOVs for each concentration. **e,f,g** Single-molecule detection of p53 in the plasma of healthy and CRC patients with known p53 mutations. **e,** Representative TIRF images. Detection is carried out simultaneously with antibodies targeting all p53 (red) and with antibodies that are specific to the mutant p53 conformation (green). Large diameter spots correspond to TetraSpeck beads used for alignment. **f,** p53 signal accumulation over time in late stage CRC and healthy plasma. Data is presented as the mean +/- s.d. of 50 FOVs for each time point. **g,** Total and mutant p53 levels in plasma show significantly higher levels in CRC patients versus healthy individuals (n=6 for each group). Box plots limits: 25–75% quantiles, middle: median, upper (lower) whisker to the largest (smallest) value no further than 1.5× interquartile range from the hinge. P values were calculated by Wilcoxon rank sum exact test. *** P value < 0.001. **h,** Experimental scheme for single-molecule imaging of global DNA methylation: MBD2-biotin is incubated with Cy3-labeled (green) DNA, and binds specifically to methylated DNA molecules. Next, biotin-MBD-meDNA complexes are immobilized on a PEG-streptavidin surface, followed by TIRF imaging. Each spot represents a single bound complex, number of spots correspond to the level of DNA methylation in plasma. **i,** Representative TIRF images of DNA methylation in HEK293 cells treated with 5-Aza-2′-deoxycytidine, demonstrating significant reduction in methylation compared to control cells. **j,** Representative TIRF images of global cfDNA methylation levels in the plasma of CRC and healthy subjects, showing lower DNA methylation levels in CRC. For all images, numbers within images represent counted spots.

Counting of single molecules confers high sensitivity^27, 28^, thus we explored whether we could also quantify non-secreted tumor-specific plasma proteins that are undetectable by conventional technologies. We focused on the tumor suppressor p53, which is frequently mutated in CRC; p53 mutations lead to its stabilization and accumulation in tumor cells^29^. We captured p53 from plasma and applied simultaneous detection with two distinct antibodies; an antibody targeting both the wild type and mutant forms of p53, or another antibody specifically targeting the mutant conformation (Fig. 2e). Time-lapse imaging enabled the accumulation of p53 signal, overcoming the transient binding dynamics of the detection antibodies (Fig. 2f and Supplementary Fig. 4d). Indeed, we observed higher levels of total and mutant p53 in the plasma of CRC patients with confirmed p53 mutations (Fig. 2g), establishing our system’s capabilities in specific detection of mutant proteins that originate directly from tumor cells.

DNA methylation is often deregulated in cancer, and specifically in colorectal cancer^30, 31^. We therefore aimed to combine our analysis with quantitative single-molecule detection of DNA methylation levels in plasma. We incubated Methyl-CpG-binding domain protein 2 (MBD2-biotin), which specifically binds to methylated DNA^32^, with fluorescently labeled plasma cfDNA. Bound complexes were anchored to the surface and imaged (Fig. 2h). Of note, bound DNA molecules may harbor one or more methylation sites. Specificity and sensitivity were validated using synthetic methylated/unmethylated DNA, as well as DNA from cells treated with the DNA methyl transferase (DNMT) inhibitor 5-Aza-2′-deoxycytidine (Fig. 2i and Supplementary Fig. 5a-c). Finally, we verified detection of cfDNA methylation levels from plasma of CRC and healthy subjects (Fig. 2j and Supplementary Fig. 5d).

We applied EPINUC to generate high-dimensional data, comprising of the three layers of information; histone PTMs, DNA methylation and protein biomarkers, from 33 plasma samples of healthy subjects and 46 samples taken from 40 late stage CRC patients (stages Ⅲ-Ⅳ; six patients were sampled twice at different times during cancer progression and treatment). CRC samples were obtained from patients prior to surgery or from patients that underwent surgical resection procedure and chemotherapy. In accordance with its use in clinical diagnostics^24^, single-molecule counting of CEA showed higher levels in CRC patients (Fig. 3a and Supplementary Fig. 6a,b), and a reduction in patients after resection. Interestingly, high CEA levels were also observed in a few healthy individuals, generating a ‘false positive’ signal (for example, samples 6, 9 and 19, marked by * in Supplementary Fig. 6a). Simultaneous probing of MST1, an anti-proliferative factor, allowed us to derive the CEA/MST1 ratio, resulting in better classification of samples and highlighting an advantage of combinatorial biomarker detection (Fig. 3b,c and Supplementary Fig. 6a). Of note, plasma from CRC patients following resection exhibited altered CEA/MST1 ratio compared to non-resected patients, showing higher similarity to healthy individuals (Fig. 3a,b). This demonstrates the potential applicability of our technology to monitor treatment, while underlying the need to collect additional information from each sample to allow correct sample classification.

**Fig.3:**
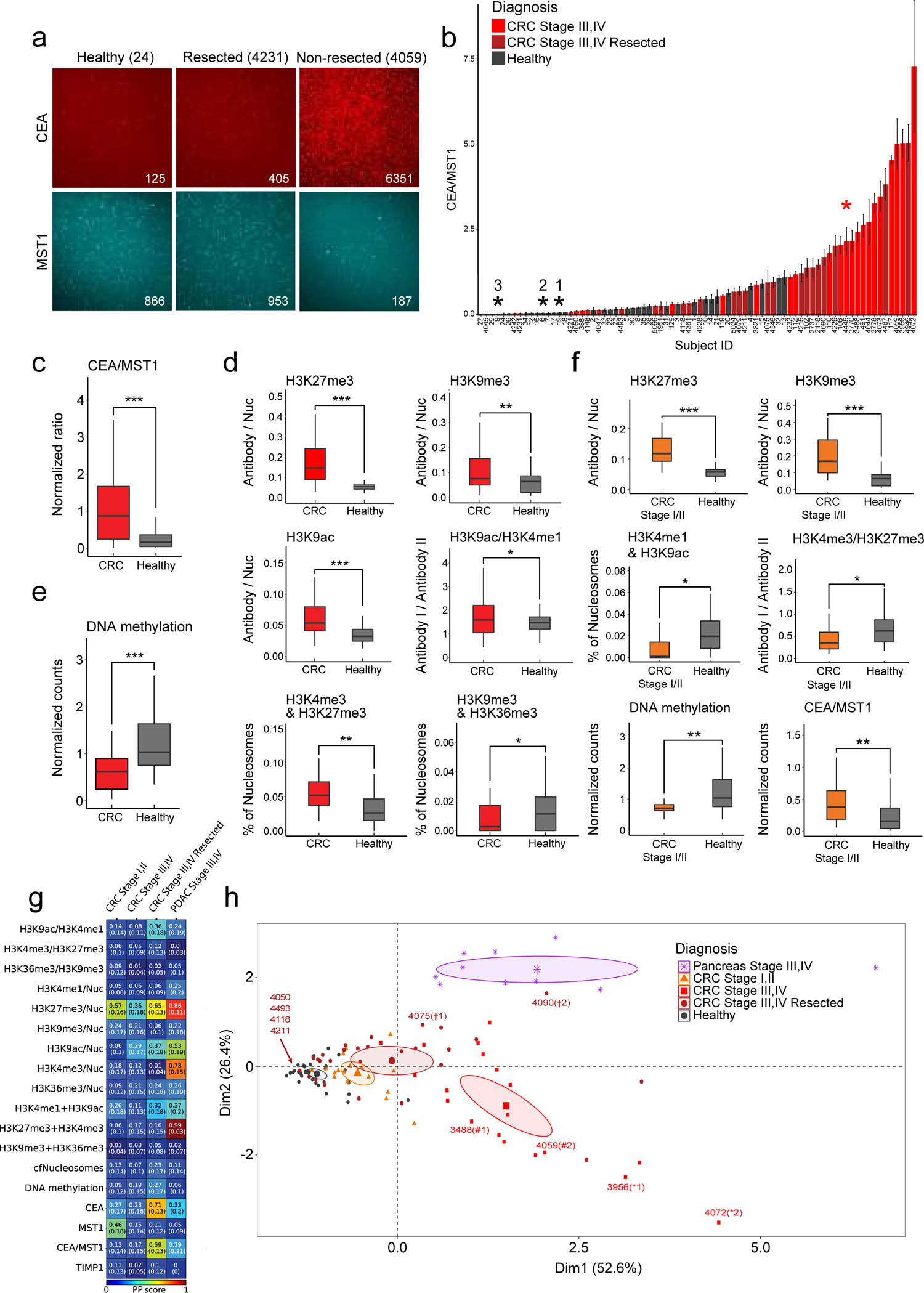
EPINUC reveals significant epigenetic and biomarkers alterations in the plasma of CRC patients. **a,** Representative TIRF images depict changes in protein biomarker levels in the plasma of healthy, CRC patient, and CRC following tumor resection. Numbers within images represent counted spots. **b,** CEA/MST1 normalized levels in the plasma of CRC patients and healthy individuals. Each bar represents a subject; data is presented as the mean +/- s.d. of 50 fields of view per sample. *1-3 correspond to healthy samples 19, 6, and 9, respectively. Sample 4445 (CRC, red) is denoted by *. **c,** Box plot representation of the data in B (healthy = 33, CRC = 46(. Box plots limits: 25–75% quantiles, middle: median, upper (lower) whisker to the largest (smallest) value no further than 1.5× interquartile range from the hinge. P values were calculated by Welch’s t-test. *** P value < 0.001. **d,** Histone PTMs, ratios and combinations (as indicated on the graphs) that significantly differ between healthy and CRC late stage samples (healthy = 33, CRC = 46(. P values were calculated by Welch’s t-test. * P value < 0.05 ** P value < 0.01. *** P value < 0.001. **e,** Global DNA methylation levels, measured as in Fig. 2J, in the same cohort as (d). **f,** EPINUC measurements (Histone PTMs, DNA methylation and protein biomarkers) that significantly differ between healthy and early stage CRC patients (healthy = 33, early CRC = 17). P values were calculated by Wilcoxon rank sum exact test. * P value < 0.05 ** P value < 0.01. *** P value < 0.001. **g,** Individual parameters predictive power score (PPS) analysis for the various subgroups (see Methods). Color scale represents PPS value. **h,** Principal Component Analysis (PCA) with input parameters of H3K27me3/Nuc, H3K4me3 & H3K27me3, CEA/MST1 and CEA. Sample groups are color-coded as indicated, each dot represents a plasma sample. Ellipse represents 95% confidence interval for the barycenter of each group.

EPINUC also provides quantitative measurements of the total number of cfNucleosomes, six histone PTMs, their pairwise combinations and ratios per plasma sample (Fig. 1). In agreement with the literature, CRC patients had higher cfNucleosomes in their plasma compared to healthy controls^33^ (Supplementary Fig. 6c). While most epigenetic parameters did not change, several showed significant differences: CRC patients had higher levels of H3K27me3-, H3K9me3-, H3K9ac- and H3K4me1-modified nucleosomes, and higher ratio of H3K9ac to H3K4me1 (Fig. 3d and Supplementary Fig. 6c,d). Interestingly, the combinatorial pattern of H3K9me3+H3K36me3-modified nucleosomes decreased in CRC, concomitant with an increase in ‘bivalent’ nucleosomes marked by H3K4me3+H3K27me3 (Fig. 3d). As bivalent chromatin is strongly implicated in many types of cancers^21, 34^, this result further confirms the diagnostic value of single-molecule quantification of combinatorial histone marks. DNA methylation was reduced in CRC samples, in agreement with previous studies^35, 36^ (Fig. 3e).

The identification of epigenetic and biomarkers alterations in late stage CRC motivated us to apply EPINUC to 17 plasma samples from individuals diagnosed with early stage CRC (stages I,II). As in the later stage, the levels of DNA methylation, CEA and CEA/MST1 ratio significantly differed in early stage cancer patients versus healthy (Fig. 3f and Supplementary Fig. 6e). Interestingly, TIMP1, whose levels did not alter between our cohort of healthy and late stage CRC, was elevated at the early stage (Supplementary Fig. 6e,f). This might be due to chemotherapy treatment administrated to most of the late stage CRC patients, which was reported to downregulate TIMP1 levels^37^. On the contrary, early stage CRC patients did not receive chemotherapy, and indeed showed elevated TIMP1, rendering it a significant biomarker only for early stage. Of note, plasma from stages I,II CRC patients also showed elevated levels of H3K27me3- and H3K9me3-modified nucleosomes, as seen in the late stage (Fig. 3f). Interestingly, we did not observe increased levels of cfNucleosomes in early stage CRC, likely due to the low tumor burden (Supplementary Fig. 6e). While the levels of H3K9ac- and H3K4me1-modified nucleosomes did not differ significantly from the healthy group, the combinatorial pattern of H3K4me1 and H3K9ac was lower in early CRC patients compared to healthy (Fig. 3f). These results indicate various epigenetic alterations already occurring at early stage CRC.

As proof-of-concept that EPINUC could be generalized and applied to study diverse types of cancers, we obtained and analyzed 10 plasma samples from patients diagnosed with pancreatic ductal adenocarcinoma (PDAC), a devastating disease with poor prognosis and rising incidence^38^. The data revealed multiple epigenetic parameters, as well as protein biomarkers, which differ significantly from healthy subjects (Supplementary Fig. 7a). Interestingly, PDAC patients showed very high levels of H3K4me3- and H3K27me3-modified nucleosomes, compared to both groups of healthy and CRC patients (Supplementary Fig. 7a,b). Moreover, the percentage of bivalent nucleosomes marked by both of these modifications was surprisingly high, clearly differentiating these samples from plasmas obtained from CRC patients. Finally, as opposed to CRC, in PDAC we did not observe a significant change in H3K9me3-modified nucleosomes. Additional parameters that were altered in CRC compared to healthy, such as the combination of H3K9me3+H3K36me3 and the ratio of H3K9ac to H3K4me1, were not affected in patients diagnosed with PDAC (Supplementary Fig. 7c). Overall, while the number of samples analyzed is limited, the data suggests cancer-type specific alterations to various epigenetic parameters. Finally, we calculated the predictive score of each parameter alone to discriminate between the healthy, the distinct groups of CRC patients, and PDAC patients (Fig. 3g, Methods). These results highlight EPINUC’s capabilities in providing multiplexed single-molecule measurements of protein biomarkers, epigenetic modifications and their combinations for cancer diagnostics.

To visualize the distribution of samples across the most significant and predictive parameters, we performed Principal Component Analysis (PCA). The PCA showed spatial separation between the groups, with the early stage CRC samples positioned in between the healthy and the late-stage CRC, potentially reflecting a transition stage (Fig. 3h). While samples from healthy individuals formed a tight cluster, the cancer samples showed greater variability, likely due to inherent heterogeneity between tumors. CRC patients who underwent resection also exhibited a high heterogeneity; interestingly, patients who received both primary tumor resection and metastectomy were positioned closer to the healthy group (samples 4118, 4211, 4050 and 4493). Strikingly, the PDAC samples clustered separately from the CRC samples, pointing to the power of EPINUC in differentiating cancer types solely based on multiplexed measurements of epigenetic parameters.

A few CRC patients in our cohort were sampled twice along the course of the study, allowing us to examine the projection of samples taken from the same individual in the PCA plot (Fig. 3h, marked in *, ^#^, and †). Sample 4090 was collected two months following sample 4075 from a CRC patient who underwent tumor resection and extensive treatments. Unfortunately, her condition did not improve and she passed away a month later; indeed the later sample projects further from healthy on both principle components. A similar trend can be seen for samples 3488 and 4059, taken 6.5 months apart. These results highlight the potential of EPINUC to monitor patients’ positive or negative response to treatment, and the power of collecting multiple layers of information from each sample.

Finally, in order to integrate all measurements and fine-tune the discrimination between healthy and CRC samples, we employed machine-learning classification (Fig. 4a, Methods). The best predictive model displayed high diagnostic potential by generating a 0.96 AUC [95% confidence interval (CI) 0.945 - 0.975], and sensitivity of 92% [95% CI 89.3 – 94.7] at 85% specificity [95% CI 80.2 – 89.8] and 92% precision [95% CI 89.7 – 94.3], outperforming predictive models relying solely on protein biomarkers, protein biomarkers coupled with DNA methylation, or protein biomarkers coupled with histone PTMs (Fig. 4a and Supplementary Fig. 8a). Intriguingly, this high discrimination power is achieved without including DNA sequencing. This is mainly due to the combination of multiple parameters spanning various cellular pathways into a single assay, and the high accuracy of the single-molecule technology that allows for digital counting of molecules.

**Fig.4:**
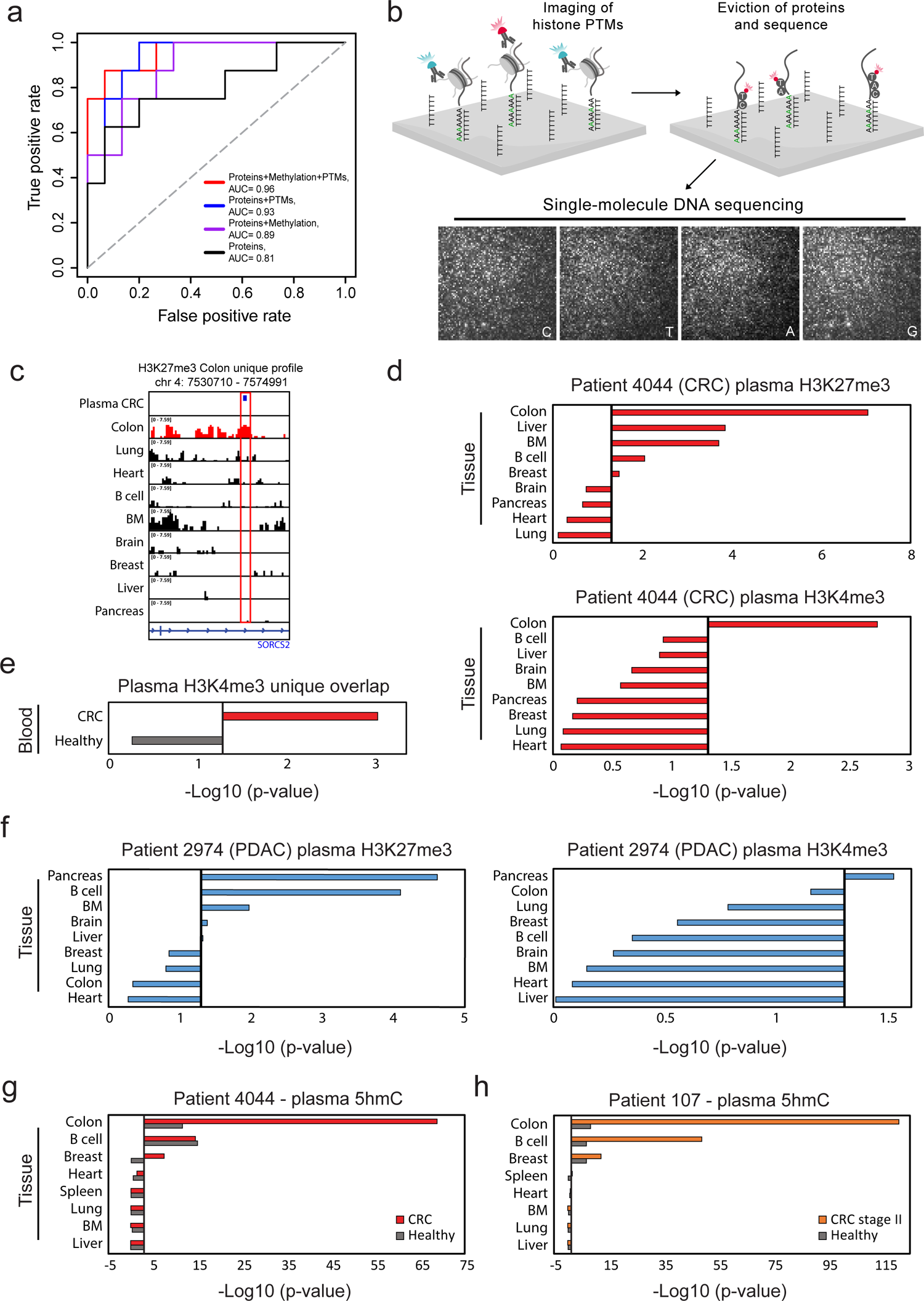
EPINUC differentiates healthy versus CRC patients and informs the tumor tissue-of-origin. **a,** ROC curves discriminate between healthy (n=33) and all CRC samples (n=63) using a logistic regression model. The area under the curve (AUC) is shown for protein biomarkers only (Proteins; CEA + CEA/MST1, black line), protein biomarkers in combination with global DNA methylation levels (Proteins + Methylation, purple line), protein biomarkers with histone PTMs (Proteins + PTMs, blue line) and the complete dataset generated by EPINUC (Proteins + Methylation + PTMs, Red line). Gray diagonal line indicates expected curve for random classification. **b,** Experimental scheme for EPINUC-seq. Histone PTMs are decoded as shown in Fig. 1A. Next, histones are evicted by increasing salt concentration, retaining DNA strands at identical positions. Single-molecule DNA sequencing-by-synthesis is performed by cycles of incorporation of fluorescently labeled nucleotides and TIRF imaging^21^. Images represent four sequencing cycles, showing incorporation of cytosine (C), thymine (T), adenosine (A) and Guanine (G). For each x,y coordinate on the surface, sequence data is analyzed and integrated with the initial images that registered histone PTMs, revealing the modification state of the corresponding nucleosome. **c,** Single-molecule reads of H3K27me3 (blue) from a CRC patient overlap with ChIP–seq profile of H3K27me3 in the colon, but not the other indicated tissues. BM=bone marrow. **d,** EPINUC-seq analysis of plasma from stage IV CRC patient (patient 4044). Tissues and primary cell lines ranked by overlap significance with single-molecule plasma H3K4me3 (top) or H3K27me3 (bottom) positive reads. Black line corresponds to P value of 0.05. P values were determined by Z-test. **e,** Overlap significance of H3K4me3 single-molecule reads with ChIP-seq data^13^ performed in the plasma of healthy and CRC patients. **f**, EPINUC-seq analysis of plasma from stage IV PDAC patient (patient 2974). Tissues and primary cell lines ranked by overlap significance with single-molecule H3K27me3 (Left) or H3K4me3 (Right) positive reads. **g,h,** Overlap significance of tissues and primary cell lines unique H3K36me3 profiles with single-molecule 5hmC reads from healthy versus late stage CRC (**g**) or early stage CRC (**h**), calculated as in (e). Black line corresponds to P value of 0.05. P values were determined by Z-test.

We hypothesized that introducing a sequencing feature for samples that were classified as cancerous by the machine-learning algorithm would provide yet another layer of specificity and sensitivity. As different tissues vary in their epigenetic modifications, it may allow detection of the tissue-of-origin of the circulating nucleosomes, thus revealing the origin of the cancer. To that end, we coupled the epigenetic analysis with single-molecule DNA sequencing^21^ (Fig. 4b and Supplementary Fig. 8b). Briefly, following detection of histone PTMs on cfNucleosomes, the histone proteins are evicted, and the DNA is subjected to repeated cycles of sequencing-by-synthesis using an automated fluidics system. Each cycle consists of incorporation of A, C, T or G by DNA polymerase and imaging; following 120 cycles, the data is integrated to build a strand that can be aligned to the genome, corresponding to the position of the modified nucleosome.

As proof-of-concept, we applied EPINUC followed by sequencing (EPINUC-seq) to three plasma samples of late stage CRC probed for H3K4me3, H3K27me3 or H3K9ac (Fig. 4c,d and Supplementary Fig. 8c-e). Single-molecule mapped reads, corresponding to modified nucleosomes, were intersected with unique antibody peak signatures generated from ENCODE ChIP-seq data for various tissues and primary cell lines, followed by bootstrapping simulations to calculate significance (Supplementary Fig. 8c, Methods). Reinforcing our hypothesis, we found that plasma samples showed significant overlap with colon-specific H3K4me3, H3K27me3 and H3K9ac peaks, indicating colon as the main tissue-of-origin (Fig. 4c,d and Supplementary Fig. 8c-e). Moreover, comparing our data to a recent ChIP-seq study of H3K4me3 in plasma showed significant overlap with profiles obtained from CRC patients^13^, but not with healthy plasma (Fig. 4e). H3K27me3 mapped reads showed a broader pattern compared to H3K4me3 and H3K9ac, overlapping with peaks corresponding to hematopoietic lineage as well as colon. Interestingly, for CRC patients 4044 and 3821 we also observed a significant overlap with liver-specific H3K27me3 and H3K9ac peaks, in agreement with clinical data indicating these patients had liver metastases. This is consistent with recent studies reporting liver damage and release of liver-specific DNA due to tumor cells metastasizing to the liver^39, 40^. Yet, the liver signal may also originate from chemotherapy-induced liver damage^41^. Validating our EPINUC-seq approach for tissue-of-origin detection, analysis of plasma samples from PDAC, lung and breast cancer patients revealed pancreas, lung and breast tissues as the main contributors, respectively (Fig. 4f and Supplementary Fig. 8f,g).

Finally, we combined a complementary single-molecule approach to identify the tumor tissue-of-origin, by single-molecule profiling of the DNA modification 5-Hydroxymethylcytosine (5hmC). 5hmC is known to play important roles in gene regulation and cancer^42, 43^. We captured 5hmC-enriched DNA from plasma as previously described^44^, followed by single-molecule DNA sequencing (Supplementary Fig. 9a, Methods). In agreement with previous reports^45^, we found 5hmC in cfDNA to be enriched at gene bodies and promoter proximal regions (Supplementary Fig. 9b), which are also known to be marked with H3K36me3. Thus, we generated unique 5hmC read signatures for healthy and CRC samples, and examined their overlap with H3K36me3 peak signatures from various tissues and primary cell lines (Methods). Similar to sequencing of the histone marks (Fig. 4d), analysis of 5hmC in the plasma of the same patient showed highly significant overlap with the colon-specific profile, validating this strategy for identification of the tumor tissue-of-origin (Fig. 4g and Supplementary Fig. 9c). We next aimed to explore whether we can correlate the 5hmC signatures with gene expression data from primary CRC tumors. Thus, we generated a list of genes found to be associated with 5hmc signal for each group of healthy and CRC patients. Interestingly, the expression levels of these CRC-specific genes (identified based on the 5hmc signal) was significantly higher in CRC tumors versus the expression levels of the group of healthy-specific genes (Supplementary Fig. 9d). Finally, we showed correct identification of colon origin also for early stage CRC patients, and identification of pancreas origin for the patient diagnosed with PDAC (Fig. 4h and Supplementary Fig. 9e,f).

Our work establishes EPINUC as a novel liquid biopsy approach that analyzes multiple histone and DNA modifications, as well as protein biomarkers, at single-molecule precision. EPINUC distinguishes between CRC patients to healthy individuals at high specificity and sensitivity. We showed that this multi-parametric approach is suitable also for detection of early stage patients, although expanding the analysis to a larger cohort is needed. The main challenges with analyzing plasma nucleosomes are (1) their minute amount-in 1 ml of plasma there are ∼1000 genome copies^13, 46^; (2) The plasma is highly dense with additional proteins, rendering enzymatic or binding approaches to capture nucleosomes difficult; (3) There is high variability between different individuals, stressing the need for quantitative methodologies to allow comparison between samples; and (4) Multi-parametric data is needed to achieve high specificity and confidence in detection. Our EPINUC approach addresses these challenges by enabling single-molecule combinatorial detection of epigenetic marks, DNA sequencing and protein biomarkers from limited input material. For quantification of global DNA methylation levels, several alternative approaches have been developed^47, 48^. The methylation status of repeat elements in the genome, measured by bisulfite conversion followed by PCR, is frequently used as proxy for global genomic methylation. Yet, it is uncertain to what extent this method provides accurate representation of global DNA methylation in diverse biological and pathological conditions. Alternative approaches utilize mass spectrometry to quantify methylated versus unmethylated cytosine; while these methods are quantitative and accurate, they require relatively high amounts of DNA. Our approach complements these methodologies by providing quantitative single-molecule measurements of global DNA methylation, which is independent on chemical conversion and is not limited to analysis of specific genomic regions, and requires very low input material.

In addition to the unique epigenetic analysis, the single-molecule system outperforms the classical ELISA assay for measuring protein biomarkers. ELISA is of relatively low sensitivity and is therefore limited to proteins that are present at high levels, has lower dynamic range in quantifying proteins, and is not amenable to multiplexed detection of several proteins^27, 28^. We showed that the single-molecule system is capable of detecting the mutant form of p53, which is a non-secreted protein that originates directly from the tumor cells. The main advantages of the single-molecule imaging system is its unique ability to follow binding of antibodies over time, thus allowing for accumulation of signal. Interestingly, the accumulation kinetics for each antibody is highly reproducible across different experiments, likely representing the antibody’s affinity and avidity to its target epitope. Importantly, the system is straightforward to adapt for detection of additional proteins, thus increasing sensitivity and enabling disease-specific biomarkers analysis.

EPINUC is built on the idea that integration of multiple parameters would provide specific diagnosis, which is independent on DNA sequencing. Indeed, we show that the few pancreatic cancer samples tested within this study cluster separately from CRC patients, despite the heterogeneity within each group. This would potentially render the EPINUC approach fast and inexpensive, paving the way to a mass screening method for a variety of cancers. Moreover, the integration of multiple parameters may overcome potential confounding effects generating heterogeneity in the cohort, such as chemotherapy treatments administrated to some of the patients, or varied composition of blood cells between individuals. Expanding the protein biomarkers panel to include cancer-type specific indicators may be instrumental in allowing differential diagnosis, independent of sequencing data. Nevertheless, further studies, analyzing larger cohorts of patients, are needed to test whether this technology can differentiate between various cancer types based solely on the epigenetic and biomarker profiles. Such studies may also provide insights on whether the epigenetic differences detected by EPINUC originate solely from the tumor cells, or may represent, at least in part, epigenetic alterations occurring in non-cancer cells in the tumor microenvironment, or generated by chemotherapy or other treatments. Importantly, we showed that the combination of EPINUC with single-molecule DNA sequencing provides unequivocal evidence of the tumor tissue-of-origin, thus further elucidating the type of cancer of each individual. In future studies, it would be of special interest to apply EPINUC-seq to cancer of unknown primary, potentially identifying its origin. Further efforts in clinical use of EPINUC would require a system that offers streamlined sample workflow, integrated instrumentation, and robust data processing pipelines and interpretation algorithms. EPINUC technology greatly expands the already burgeoning field of liquid biopsies and has the potential to be applicable for early detection of cancer and monitoring.

## Acknowledgments

We thank Dr. R. Rozenzweing, Dr. O. Fasust, Mr. M. Maurer, and Ms. R. Irwin for their contribution in establishing a protocol for Methyl-CpG-binding domain protein 2 labeling. We thank Lior Segev for his help with writing and integrating the μManager scripts for performing EPINUC-Seq. We are grateful for the important comments made by I. Ulitsky while reading the manuscript.

## Author contributions

V.F, N.E, and E.S designed the study and wrote the manuscript. V.F and N.E performed the experiments and analyzed the data. N.F and O.B assisted in the experiments, G.R assisted with data analysis. B.B.Z, Y.M, T.P, A.H, J.E.C, A.S, M.T, A.G, M.M and A.Z collected the plasma samples of early and late stage CRC patients. D.J, A.S, K.A contributed to the development of single-nucleosomes imaging technology and sequencing experiments described in this study.

## Data and materials availability

All data is available in the main text or the supplementary materials.

## Methods

### Patients

All clinical studies were approved by the local ethics committees (Helsinki applications 091-2020 and 0198-14-HMO). Informed consent was obtained from all individuals before blood sampling.

### Plasma collection

Blood samples were collected in VACUETTE K3 EDTA tubes and transferred immediately to ice. Next, blood was centrifuged (10 minutes, 1,500g, 4°C) and the supernatant was transferred to a fresh 50-ml tubes and centrifuged again (10 minutes, 3,000g, 4°C). The supernatant was collected and used as plasma for all experiments. Plasma was analyzed fresh or flash-frozen and stored at −80°C for future analysis.

### Cell-free nucleosomes (cfNucleosomes) preparation for single-molecule imaging

Tagging and tailing of cfNucleosomes was carried out as following: 20 µ l of plasma or 5X (DDW diluted) concentrated apoptotic medium was incubated at 37°C for 1 hour with the following reaction mixture: 10 µ l 10X Green Buffer (Enzymatics, B0120), 416 µ M CoCl2 (Enzymatics, B0220), 1:60 PI (SIGMA, P8340), 83.3 nM fluorescently labeled dATP (Jena Bioscience, NU-1611-Cy3/Cy5), 83.3 µ M dATP (Thermo Fisher Scientific, R0181), 5 µ l of Klenow Fragment (3’→5’ exo-, NEB, M0212S) 3µl of T4 Polynucleotide Kinase (NEB, M0201L) and 4 µ l of Terminal deoxynucleotidyl transferase (TdT, Enzymatics, P7070L). Following incubation, samples were inactivated by immediate transfer to ice. For nucleosome sequencing, 1.67 µ l of ddATP was added (SIGMA, GE27-2051-01).

### Plasma cell-free DNA (cfDNA) isolation and fluorescent labeling

cfDNA was extracted from 4 ml of healthy human blood plasma, or from 0.5 ml of plasma from CRC patients, using the Mag-Bind cfDNA Kit (Omega Bio-Tek, M3298-01). For optimized yield, protocol was modified by increasing elution time to 20 minutes on a thermomixer, at 1,600 rpm, in 15 µl elution buffer at room temperature. Sample concentration was measured using Qubit Fluorometer (Thermo Fisher Scientific). For fluorescent labeling of plasma isolated DNA, 10 µ l of cfDNA was incubated at 37°C for 1 hour with the following reaction mixture: NEBuffer™ 2 (NEB, B7202), 0.25 mM MnCl2 (SIGMA, M1787), 33 µM fluorescently labeled dATP (Jena Bioscience, NU-1611-Cy3), 1.5 µ l of Klenow Fragment (3’→5’ exo-, NEB, M0212S) and 1.5 µ l of T4 Polynucleotide Kinase (NEB, M0201L). Following incubation, samples were inactivated by addition of EDTA (Invitrogen, 15575-038) at a final concentration of 20mM. Next, DNA was purified by AMPure SPRI beads (Beckman Coulter, A63881), and quantified by Qubit (Thermo Fisher Scientific).

### Cell culture and apoptosis

Cell lines were maintained at 37°C with 5% CO2. HEK-293 cells were cultured in 150 cm plates (10×10^6^ cells in 20 ml of media) in DMEM supplemented with 10% FBS and 1% P/S, and passaged every week. For induction of apoptosis, 6 µ M of Staurosporine (STS, Holland-Moran, 62996-74-1.25) was added to medium of confluent cells. 72 hours later, medium was collected and immediately processed. To verify fragment sizes along with nucleosome labeling, 10 ul of the nucleosomes and 10ul of AMPure extracted DNA (either directly from concentrated medium or after the tagging and tailing reaction), were loaded on High Sensitivity D1000 ScreenTapes (Agilent, 5067-5584) and 6% TBE gel (ThermoFisher Scientific, EC62655BOX), and imaged with 4200 TapeStation (Agilent) or Typhoon imager (Amersham Biosciences), respectively. Apoptotic medium cfNucleosomes were concentrated and recovered using Centricon Plus-70 centrifugation filteres (Merck, UFC710008) according to the manufacture protocol. PI was supplemented 1:100 following concentration.

### Surface preparation for single-molecule imaging

PEGylated-Biotin and PEGylated-poly T coated microscope slides were prepared based on the protocol described by Chandradoss et al^49^. Ibidi glass coverslips (25 mm x 75 mm, IBIDI, IBD-10812) were cleaned with (1) MilliQ H2O (3X washes, 5 minutes sonication, 3X washes); (2) 2% Alconox (SIGMA, 242985) (20 min sonication followed by 5X washes with MilliQ H2O); and (3) 100% Acetone (20 min sonication followed by 3X washes with MilliQ H2O). Slides were further cleaned and functionalized (Hydroxylated) by incubation in 1 M KOH (SIGMA, 484016) solution for 20 minutes while sonicated, followed by 3X washes with MilliQ H2O. Slides were sonicated twice for 10 minutes in 100% HPLC ethanol (J.T baker 8462-25) prior to applying amino-silanization chemistry. Next, slides were incubated for 24 minutes in a mixture of 3% 3-Aminopropyltriethoxysilane (ACROS Organics, 430941000) and 5% acetic acid in HPLC EtOH, with 1 minute sonication in the middle. Slides were then washed with HPLC EtOH (3X) and MilliQ H2O (3X) and dried with N2. Surface functionalization along with first passivation step was performed by applying mPEG: PEGylated-Biotin/PEG-Azide solution [20 mg PEGylated-Biotin (Laysan, Biotin-PEG-SVA-5000), 180 mg mPEG (Laysan, MPEG-SVA-5000) or 20 mg PEG-Azide (JenKem, A5088), 180 mg mPEG (Laysan, MPEG-SVA-5000)] dissolved in 1560 ul 0.1 M Sodium Bicarbonate (SIGMA, S6297) and degassed (centrifugation at 1 minute at 16,000*g*). Next, 140 µ l of solution was applied on one surface, followed by immediate assembly of another surface on top. Each pair of assembled surfaces were incubated overnight in a dark humid environment.

For PEGylated-Biotin surfaces: At the next day, surfaces were washed with MilliQ H2O and dried with N2 followed by a second passivation step. MS(PEG)4 (ThermoFisher Scientific, TS-22341) was diluted in 0.1 M of sodium bicarbonate to a final concentration of 11.7 mg/ml and applied on one surface, followed by the assembly of another surface on top. Each pair of assembled surfaces were incubated overnight in dark humid environment. The next day, surfaces were disassembled, washed with MilliQ H2O and dried with nitrogen. After nitrogen flush, surfaces were stored in −20°C.

For PEGylated-poly T surfaces, following PEG-Azide coating, surfaces were washed with MilliQ H2O and dried with N2. To enable anchoring of dT50 to surface via click chemistry, 10 µM of 5’heyxynyl-dT50 (IDT) were mixed with 2 mM of CuSO4 (SIGMA, C1297) and DDW. Next, 100 µ l of the mixture was applied on one surface, followed by immediate assembly of another surface on top. Each pair of assembled surfaces was incubated overnight in a dark humid environment. In the next day, a second passivation step [MS(PEG)4] was carried out, similarly to PEGylated-Biotin preparation. Surfaces were stored in −20°C post nitrogen flush in a similar fashion.

### Antibody labeling

Capture and detection antibodies were labeled using Biotin conjugation kit (Abcam, ab201796) and Alexa flour antibody labeling kits (Thermo Fisher Scientific, A20181/ A10237/A20186) according to the manufacture protocol. Labeled antibodies were purified by size exclusion chromatography using Bio-Spin 6 columns (Bio-Rad, 7326200) followed by measurement of protein concertation using *Nanodrop* 2000 at 260 nm.

### TIMP-1 siRNA transfections

siRNA transfection was performed using INTERFERin (Polyplus, 409-10) according to the manufacturer’s protocol. Briefly, cells were plated in 6-well plates (1.5 × 10^5^ in 2.5 ml per well) overnight, and the 200 µ l of transfection complex was added directly to medium, at final concentration of 25 nM of siRNA. RNA and protein samples were isolated from cells 72 hours after transfection. The following siRNA was used: SMARTpool: ON-TARGETplus Human TIMP1 siRNA (L-011792-00-0005, Dharmacon). For single-molecule imaging, medium was collected from plates, followed by centrifugation at max speed in 4°C and collection of supernatant to separate proteins from cell debris. Protein concentration was determined by Pierce™ BCA Protein Assay (Thermo Fisher Scientific, 23225), followed by addition (1:100) of protease inhibitor cocktail (PI, SIGMA, P8340).

### Synthetic DNA preparation for DNA methylation assay

DNA fragments were generated by conventional PCR (primer sites underlined) supplementing the reaction with either methylated (NEB, N0356S) or un-methylated cytosine (Thermo Fisher Scientific, R0181), followed by purification with AMPure SPRI beads. The size (∼200 bp) was chosen to mimic the size of mono-nucleosomal DNA fragments previously identified in blood plasma^50^. Fragment labeling, purification and quantification was performed as described for plasma cfDNA.

Sequence: <u>CATCAATGTATCTTATCATGTCTG</u>TATACCGTCGACCTCTAGCTAGAGCTTGGCGT AATCATGGTCATAGCTGTTTCCTGTGTGAAATTGTTATCCGCTCACAATTCCACAC AACATACGAGCCGGAAGCATAAAGTGTAAAGCCTGGGGTGC<u>CTAATGAGTGAGC TAACTCACA</u>

### 5-Aza-2-Deoxycytidine Treatment

HEK-293 cells were plated in 150 cm plates (10×10^6^ cells in 20 ml of media) overnight, then treated with 1 uM of 5-Aza-2-deoxycytidine (5 –Aza, SIGMA, A3656) or PBS for 4 days. Next, 5×10^6^ cells were collected and washed with PBS supplemented with PI (1:100), followed by centrifugation at 3000 rpm for 3 minutes. Cell pellet was resuspended with 1 ml of 0.05% Igepal (SIGMA, I8896) diluted in PBS (supplemented with PI as mentioned above) and centrifuged again at 3000 rpm for 3 minutes. Next, the pellet was resuspended in Lysis buffer [100 mM Tris-HCl pH 7.5 (Gibco, 115567-027), 300 mM NaCl (J.T Baker, 7647145), 2% Triton® X-100 (SIGMA, 9002 93-1), 0.2% sodium deoxycholate (SIGMA, D6750), 10 mM CaCl2 (SIGMA, 21115)] supplemented with PI and Micrococcal Nuclease (ThermoFisher Scientific, 88216). The reaction mixture was incubated at 37°C for 10 minutes and then inactivated by addition of EGTA at a final concentration of 20mM. Then, lysate was centrifuged for 10 minutes at max speed and supernatant was transferred to a new tube. DNA extraction, fluorescent labeling and quantification was performed as described for plasma cfDNA.

### Single-molecule imaging

PEGylated-Biotin and PEGylated-poly T coated coverslips were assembled into an Ibidi flowcell (Sticky Slide VI hydrophobic, IBIDI, IBD-80608) generating a six lane flowcell, which enables imaging of six different samples or various combinations of antibodies on a single surface. For PEGylated-Biotin flowcells, Streptavidin (SIGMA, S4762) was added to a final concentration of 0.2 mg/ml followed by 10 minutes incubation and washing with imaging buffer [IMB: 12 mM HEPES pH 8 (Thermo Fisher Scientific, 15630056), 40 mM TRIS pH 7.5 (Gibco, 115567-027) 60 mM KCL (SIGMA, 60142), 0.32 mM EDTA (Invitrogen, 15575-038), 3 mM MgCl2 (SIGMA, 63069), 10% glycerol (Bio-Lab, 56815), 0.1 mg/ml BSA (SIGMA, A7906) and 0.02% Igepal (SIGMA, I8896)]. For time-lapse imaging experiments (Histone PTMs, p53), prior to sample application, TetraSpeck beads (ThermoFisher Scientific, T7279) diluted in PBS were added and incubated on surface for at least 10 minutes to allow correction for stage drift in image analysis. Imaging was performed on a total internal reflection (TIRF) microscope manufactured by Nikon (ECLIPSE Ti2-E LU-N4 TIRF) with CFI Apochromat TIRF 60X Objective lens and TRF49904, TRF49909, and TRF89902 filter cubes (CHROMA) for the 488, 561, and 647 lasers, respectively. Images were taken with 1.5X magnification setting resulting in FOV of 148×148um, using ANDOR ZYLA 4.2 PLUS camera. At least 50 FOVs were imaged per lane.

### Histone PTMs analysis

PEGylated-poly T coated coverslips were assembled as described and further passivated with 5% BSA (Merck, A7906) for 30 minutes followed by wash with IMB. Next, plasma sample containing tailed and fluorescently labeled cfNucleosomes was incubated with antibodies (diluted 1:60) for 30 minutes at room temperature (RT), to allow formation of antibody-cfNucleosomes complexes. Next, samples were loaded on the surface and incubated for 15 minutes to allow hybridization. Flowcell was washed (X3) with IMB, followed by time lapse imaging every 15 minutes, with the three laser channels, across all positions (50 Fields of View (FOVs, 148µm^2^) per experiment). Removing antibodies from the surface (to achieve multiplexed PTM profiling) can be achieved either by multiple washes, or simply by using a reducing agent (TCEP) that disrupts the disulfide bonds between the heavy and light chains of antibodies but maintains nucleosomes intact.

### Protein biomarkers analysis

Single-molecule analysis of protein biomarkers requires the use of different capture and detection antibodies that would bind to different epitopes on the target protein. This would allow capture of target protein to the surface, while exposing a different epitope for the binding of the detection antibody. Thus, the methodology relies on the availability of two distinct antibodies for each target protein. The specific antibodies we used for each protein biomarker are listed in Supplementary Table 2.

PEGylated-Biotin coated coverslips were assembled and coated with streptavidin. Biotinylated antibodies were incubated on surface in IMB2 [10 mM MES pH 6.5 (Boston Bioproducts Inc, NC9904354), 60 mM KCL, 0.32 mM EDTA, 3 mM MgCl2, 10% glycerol, 0.1 mg/ml BSA and 0.02% Igepal] for 30 minutes, followed by wash with IMB2. Next, plasma sample was added to flowcell and incubated on surface for 30 minutes, followed by washes (3X) with IMB2, to allow binding of target proteins. Fluorescently labeled antibodies (detection antibodies) were introduced to the surface for 60 minutes, washed with IMB2, and imaged.

### Global DNA methylation analysis

PEGylated-Biotin coated coverslips were assembled and coated with streptavidin. 2 µ l of MBD2-Biotin (Thermo Fisher Scientific, A11148) was incubated with 8 µ l of Cy3 labeled cfDNA fragments for 30 minutes, to allow MBD2-Biotin binding to methylated DNA. Next, the reaction mixture was immobilized on the surface and incubated for 10 minutes, followed by TIRF imaging.

### DNA Hydroxymethylation analysis

cfDNA was incubated in 25 μl reaction mixture containing 50 mM HEPES buffer (pH 8), 25 mM MgCl_2_, 60 μM UDP-6-N_3_-Glc (Jena Bioscience, CLK-076Motif) and 12.5 U T4 beta-glucosyltransferase (Thermo Fisher Scientific, EO0831) for 2 hours at 37°C. Next, 5 μl DBCO-S-S-biotin (Click Chemistry Tools, 10 mM stock in DMSO) was directly added to the reaction mixture and incubated overnight at 37°C. DNA was cleaned using Oligo Clean & Concentrator (Zymo, D4060), and immobilized on a PEGylated-Biotin streptavidin coated surface, followed by imaging.

### Single-molecule DNA sequencing

For single-molecule DNA sequencing of cfNucleosomes, PEGylated-poly T surface was blocked with BSA as described above. Poly-A tailed FluoSpheres (described below) along with TetraSpeck beads and cfNucleosomes were applied to the surface. PTMs of plasma cfNucleosomes were imaged over time for 169 FOV, as described above. Then flowcell was washed with Wash A buffer [150 mM HEPES (KOH, pH 7.0), 1×SSC, 0.1% SDS] and Wash B buffer [150 mM HEPES (KOH, pH 7.0), 150 mM NaCl] to evict histones and antibodies. During single-molecule sequencing, temperature was maintained at 37°C by UNO-T stage top incubator (Okolab), and Luer adapter was used to connect the flowcell to IDEX 1/4-28 flat bottom fittings connecting to 1/16” OD tubing. Sequencing was performed as described previously, using Helicos True Single Molecule Sequencing (tSMS) (http://seqll.com/)^21,51^, by repurposing a microfluidics system used in the HeliScope single molecule sequencer to deliver tSMS chemistry to the flowcell)Supplementary Fig. 10). A similar setup was previously used to enable single molecule sequencing^52^.

The microfluidics system features the storage compartment for tSMS chemistry reagents, connected to a set of syringe pumps and mixing valves that deliver tSMS chemistry to the attached flowcell. During sequencing, imaging on microscope and chemistry on microfluidics system was automatically controlled by using μManager (https://micro-manager.org/) software with custom scripts to enable tSMS sequencing-by-synthesis method.

FluoSpheres preparation: FluoSpheres (Carboxylate-Modified Microspheres, Thermo Fisher Scientific, F8789) were conjugated to dA50-amine (IDT), tailed as previously described, and hybridized to the surface to serve as reference points for stage drift correction during alignment of sequencing images.

### Single-Molecule Hydroxymethylation sequencing

2.5 ng of plasma cfDNA was added to a 25 μl solution containing 50 mM HEPES buffer (pH 8), 25 mM MgCl_2_, 60 μM UDP-6-N_3_-Glc (Jena Bioscience, CLK-076) and 12.5 U T4 beta-glucosyltransferase (Thermo Fisher Scientific, EO0831), and incubated for 2 hours at 37°C. Next, 5 μl DBCO-S-S-biotin (Click Chemistry Tools, 10 mM stock in DMSO) was directly added to the reaction mixture and incubated overnight at 37°C. For 5hmC DNA pulldown, samples were incubated at RT with 15 μl of Streptavidin beads (Thermo Fisher Scientific, Dy-11205D) for 1 hour, followed by 3 washes with 1X wash buffer, and elution in 20 μl of 125 mM TCEP (Thermo Fisher Scientific, TS-77720). 5hmC eluted DNA was poly-adenylated and sequenced on a PEGylated-poly T surface as described above. Metagene profile was generated using ngs.plot.

### Image analysis

Image analysis was performed with the open-source software Cell Profiler (http://www.cellprofiler.org/). Image analysis pipelines are available upon request. Briefly, time-lapse images of antibody binding events and TetraSpeck beads are aligned, stacked and summed to one image. Antibody spots can be differentiated from TetraSpeck beads spots based on spot size and intensity. Summed antibodies images are aligned with cfNucleosomes images to count colocalization events.

### Predictive Power Score (PPS)

PPS analysis on the data was conducted using a previously published algorithm (https://github.com/8080labs/ppscore). Briefly, by calculating a cross-validated decisions tree (repeated [10K] 4-fold cross-validation) for the target variable (e.g., diagnosis) using only one of the markers, it is possible to determine which of the markers in the datasets contributes most to the target variable. The PPS is normalized to the most common assignment in order to provide a baseline for comparison. Using the PPS rather than a simple correlation measure allows us to account for non-linear effects and provides an alternative formulation for correlation which also treats categorical variables (e.g., diagnosis, or disease state – see Supplementary Table 3).

### Machine learning model for sample classification

For sample binary classification, various machine-learning algorithms were trained on the features that showed significant differences between healthy and CRC (Fig. 3c,d,e, Supplementary Fig. 6b,c), and evaluated for their performance using a four-fold cross-validation across all samples. The best predictive performance was achieved by a Logistic Regression classifier. To improve classifier performance, we conducted additional feature selection by training the classifier on all possible feature combinations out of the significant features aforementioned. Evaluating the resulting Area Under the Curve (AUC) values of repeated (500 iterations) four-fold cross-validation for each combination revealed an optimal cumulative performance of a five feature combination: H3K27me3/Nuc, H3K9me3/Nuc, CEA, CEA/MST1, and global DNA methylation. To evaluate the classifier overall performance using the selected features, we performed repeated (10K) 4-fold cross-validation across all samples. For each iteration the sensitivity, specificity, accuracy, precision, negative predictive value and the AUC value were calculated and averaged over all iterations. R caret and RWeka packages were used for machine-learning modeling.

### Tissue and plasma signatures

We downloaded and combined two independent *Homo sapiens* based ChIP-seq tracks for each tissue from the Encyclopedia of DNA Elements (ENCODE, **Supplementary Table 4**). To generate a unique antibody peak profile for a given tissue, we discarded peaks found to overlap with at least one of the other eight tissues tested, retaining only tissue specific peaks. Of note, brain H3K36me3 peaks were available only for embryonic tissue, therefore were replaced with spleen tissue H3K36me3 peaks. Similarly, heart H3K9ac peaks were available only for embryonic tissue, therefore were replaced with skeletal muscle tissue H3K9ac peaks. To generate unique plasma H3K4me3 ChIP-seq peaks, we obtained data from healthy (H) and CRC (C) plasma (n=3 for each) produced by Sadeh et. al^13^ (Supplementary Table 4). For each group, reads were intersected and only shared reads across all samples were kept for further analysis. Non-overlapping reads between the overlapping healthy and overlapping CRC reads were defined as the unique plasma signature.

### Bootstrapping simulation to analyze single-molecule reads overlap with various tissues

Overlap significance was assessed as following: First, single-molecule sequenced plasma antibody aligned reads were extended by 100bp from each side to resemble nucleosome length. Then, for each chromosome, we randomly selected a number of 230bp-long DNA segments that is equivalent to the number of antibody positive plasma reads for this chromosome. Random reads were intersected with each unique tissue/plasma signature and overlapping events were recorded. These bootstrapping simulations were iterated 10K times for each tissue for a given antibody to generate a distribution of overlap by chance. Finally, single-molecule sequenced plasma antibody reads were intersected with all tissue signatures, and contrasted against the corresponding distribution of random overlap for that tissue to evaluate overlap significance (two tailed z-test or Wilcoxon rank sum test). Signature and overlap analysis were performed using an in-house R script (EPINUC-overlap), where minimal overlap was defined as 1bp overlap. The accession numbers for the ENCODE chip-seq datasets are summarized in **Supplementary Table 4**.

### 5hmC gene signature expression in CRC

To explore the correlation between 5hmC signatures and gene expression in CRC, we generated annotated gene lists for healthy and CRC plasma (n=3 for each) enriched for 5hmC DNA sequenced reads via Clusterpofiler R package. Next, for each group, genes were intersected and only shared genes across all samples within each group were kept for further analysis. Then, in order to generate unique 5hmC gene signature for each group, both datasets were intersected and overlapping genes were discarded. Each unique 5hmC gene signature was intersected with RNA expression datasets of CRC primary tumors, obtained from the cBioportal database (TCGA, Firehose Legacy). Finally, both unique 5hmC gene signatures were compared by their logarithmic mean expression levels (two samples, Welch’s t-test).

### Statistical analysis

All statistical analysis was conducted using the statistical programming language R. Multiple comparison (Supplementary Fig. 6d) was calculated using Asymptotic K-Sample Brown-Mood Median Test.

### Data availability

Datasets generated and analyzed during this study are summarized in **Supplementary Table 4**, BED files of plasma sequenced reads are available upon request.

### Code availability

Code for performing overlap analysis is available at https://github.com/Vadim-Fed/EPINUC-overlap.

---

**Supplementary Fig.1.**
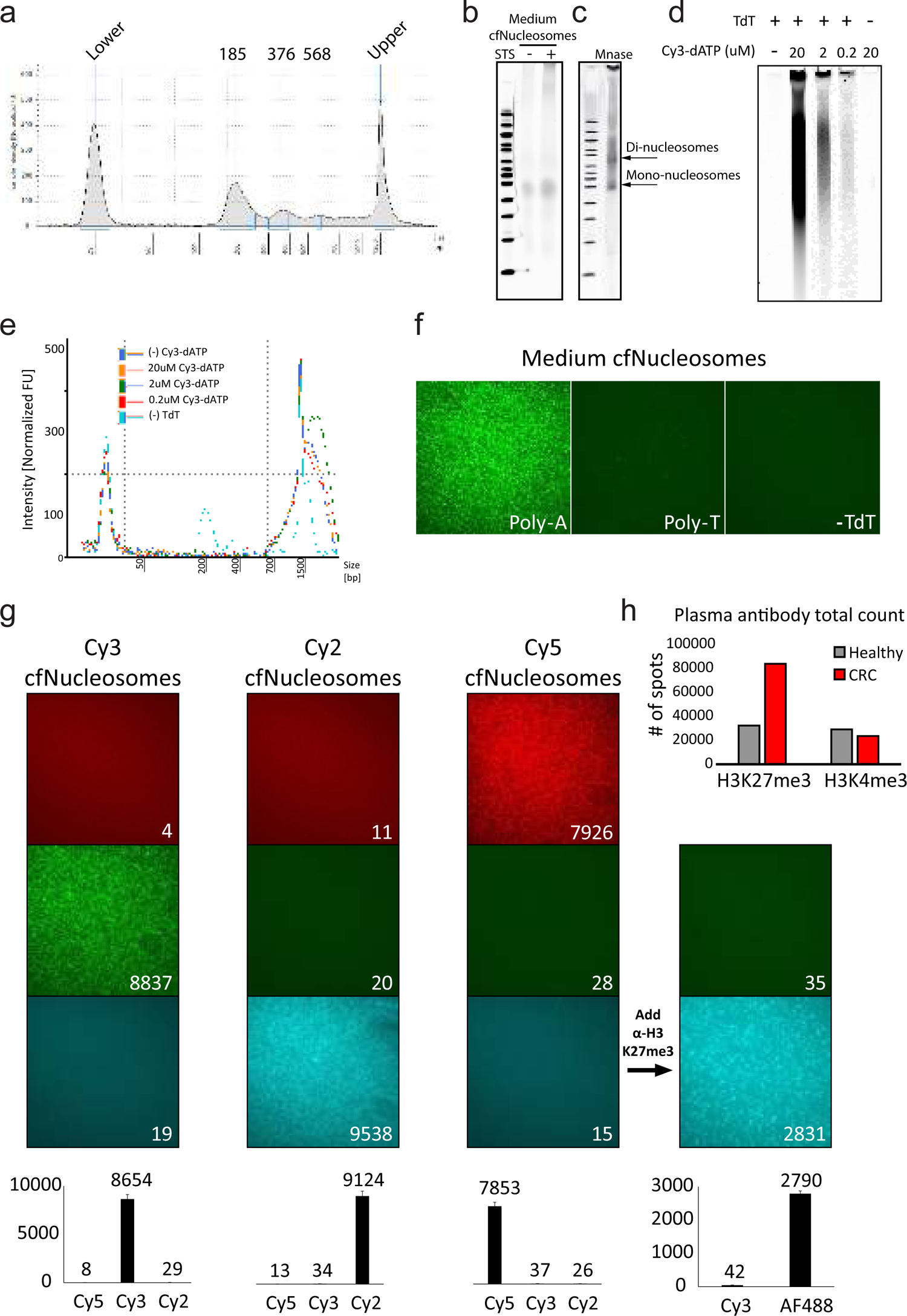
In-vitro system for sample preparation of cfNucleosomes. **(a)** TapeStation fragment size analysis of DNA isolated from medium of HEK293 cells treated with the apoptosis-inducing factor Staurosporine (STS). DNA exhibits canonical apoptotic DNA fragmentation pattern. **(b)** Nucleosomes from medium of HEK293 cells (described in a) treated with STS (+) or with PBS (-) were resolved on a 6% TBE gel and visualized by Typhoon laser scanner. **(c)** Nucleosomes were extracted from HEK293 cells by digestion with MNase^21^, resolved on a 6% TBE gel and visualized by Typhoon laser scanner. Bands correspond to mono-nucleosomes and di-nucleosomes. **(d)** Nucleosomes from medium of HEK293 cells (described in a) were subjected to tagging and tailing reaction (see Methods), resolved on a 6% TBE gel and visualized by Typhoon laser scanner. Image depicts fluorescent labeling via incorporation of Cy3-dATP during polyadenylation reaction. Incorporation is dependent on the presence of TdT in the reaction, and level of florescent signal correlates with the concentration of Cy3-dATP (Left). **(e)** TapeStation fragment size analysis of the samples in (d). **(f)** Poly-dA tailed and Cy3-dATP labeled cfNucleosomes show specific anchoring to PEGylated poly-dT surfaces through A:T hybridization. **(g)** Images and quantification of poly-dA tailed cfNucleosomes, labeled with either Cy2-dATP, Cy3-dATP or Cy5-dATP. Data indicates very low crosstalk between the three fluorescent channels. Further incubation of Cy-5-labeled cfNucleosomes with α-H3K27me3-AF488 antibody reinforces low spectral overlap. Data is presented as the mean +/- s.d. of 50 FOVs for each channel. **(h)** Total counts for H3K27me3 and H3K4me3 in the plasma samples analyzed in Fig. 1d,e.

**Supplementary Fig.2.**
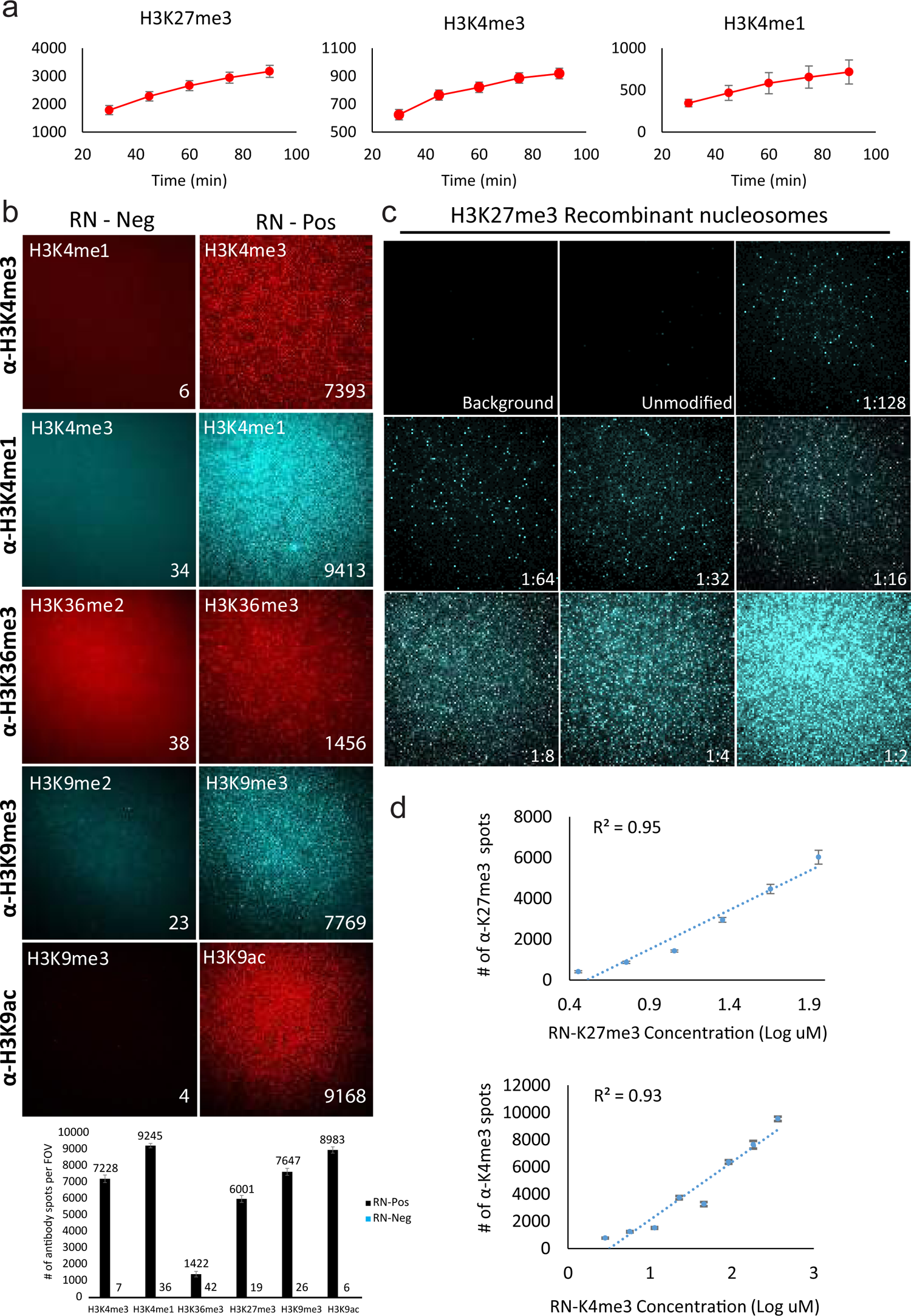
Single-molecule measurements of antibodies’ specificity and dynamics. **(a)** Accumulation of unique antibody binding events over time. Data is presented as the mean +/- standard deviation (s.d.) of 50 FOVs for each time point. **(b)** Top: Representative TIRF images demonstrate antibodies binding to various recombinant nucleosomes (RN) carrying different histone PTMs. Left images: negative control, showing very low un-specific binding of fluorescent antibodies to recombinant nucleosomes that do not carry the target PTM. Right: binding of antibodies to recombinant nucleosomes that carry the target modification. The type of modification is indicated at the bottom of the images. Numbers within images represent counted spots. Bottom: Quantification of the averaged counts of the indicated antibodies per 50 FOVs. **(c)** Representative TIRF images depicting binding of the antibody targeting H3K27me3 to an empty surface (Background), unmodified recombinant nucleosomes (Unmodified), and H3K27me3-modified recombinant nucleosomes diluted as indicated. **(d)** Regression analysis of number of spots per FOV as a function of target recombinant nucleosomes’ concentration demonstrates low variability along the regression line, supporting linearity of binding. 1:2 dilution was excluded from analysis due to high density. Data is presented as the mean +/- s.d. of 50 FOVs for each concentration.

**Supplementary Fig.3.**
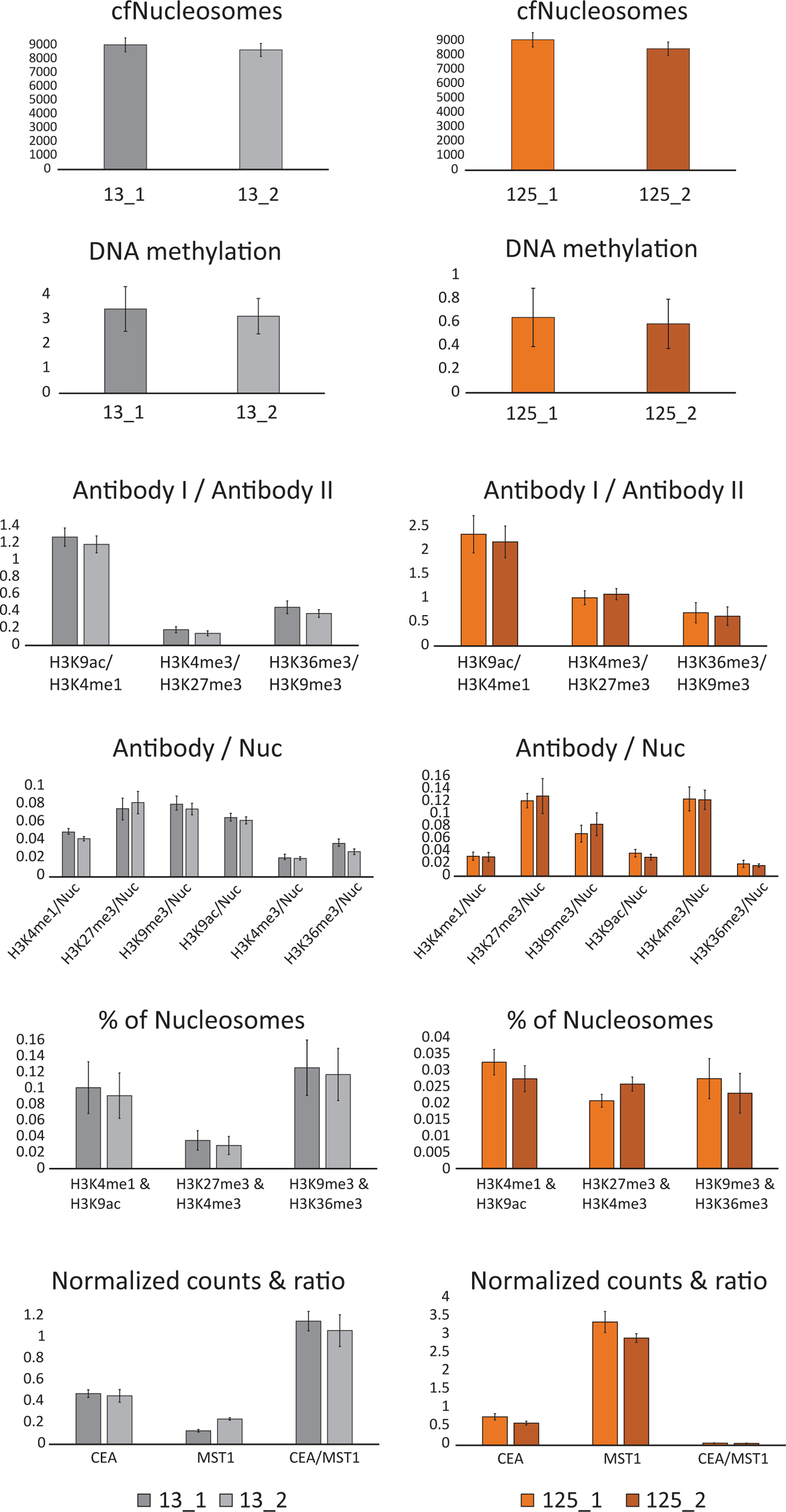
Reproducibility of EPINUC measurements. Technical repeats (n=2) of all EPINUC measurements were conducted for healthy (subject 13) and early stage (subject 125) plasma samples. Results indicate high reproducibility of EPINUC’s measured parameters, with low variation between repetitions. Of note, these technical repeats were carried out six months apart, using different batches of surfaces and different aliquots of samples. Data is presented as the mean +/- s.d. of 50 FOV for each repetition.

**Supplementary Fig.4.**
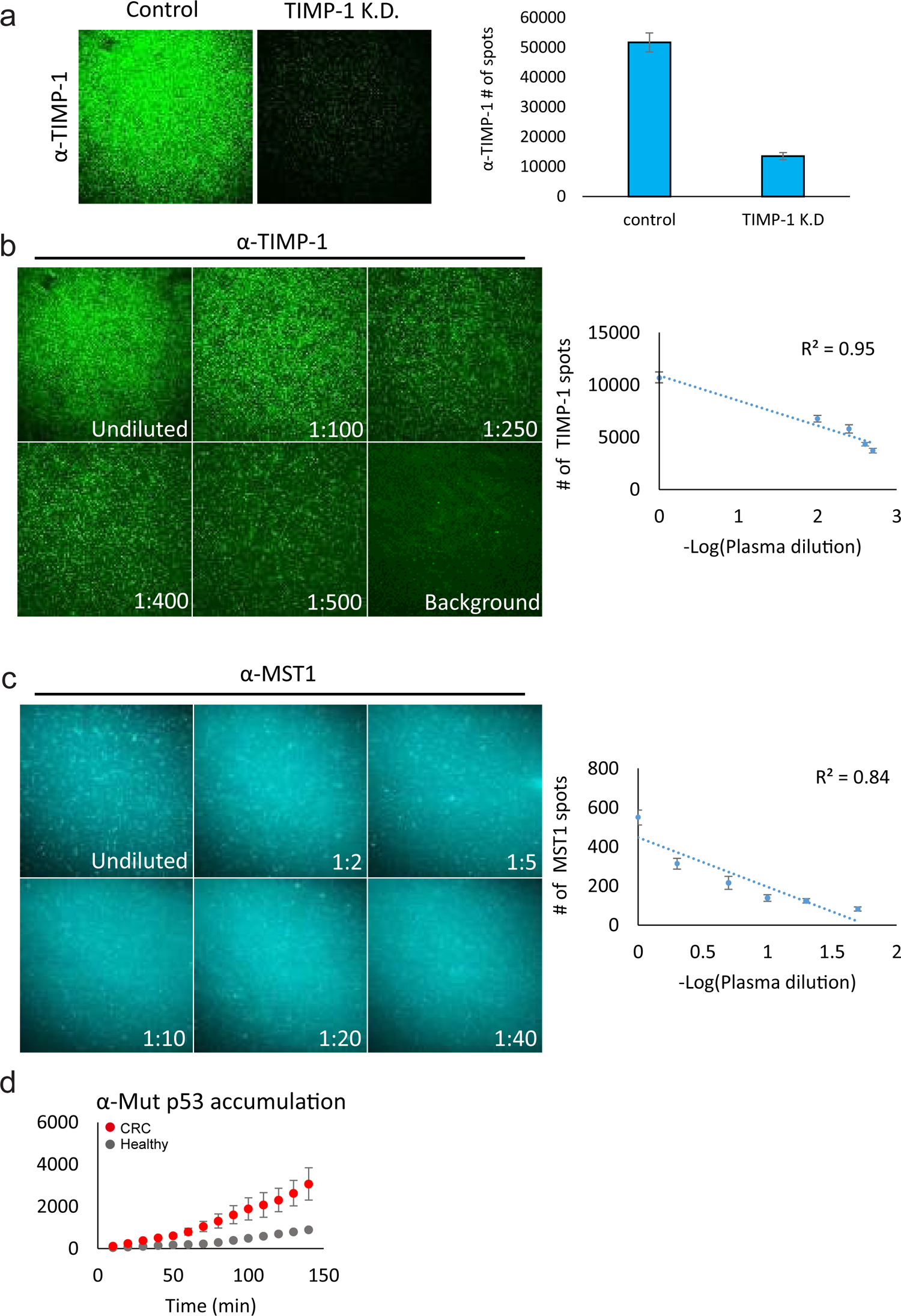
Single-molecule imaging of MST1, TIMP-1 and mutant p53. **(a)** Representative TIRF images (Left) and quantification (Right) of TIMP-1 protein levels in SW480 medium, following TIMP-1 knockdown versus control cells. Data is presented as the mean +/- s.d. of 50 FOVs for each treatment. **(b, c)** Representative TIRF images and standard curves of antibodies targeting MST1 and TIMP-1 on serial plasma dilutions, depicting linear detection of molecules within this concentration range. Data is presented as the mean +/- s.d. of 50 FOVs for each concentration. **(d)** Signal accumulation of mutant p53 over time for late stage CRC and healthy plasma samples. Each time point is presented as the mean +/- s.d. of 50 FOVs.

**Supplementary Fig.5.**
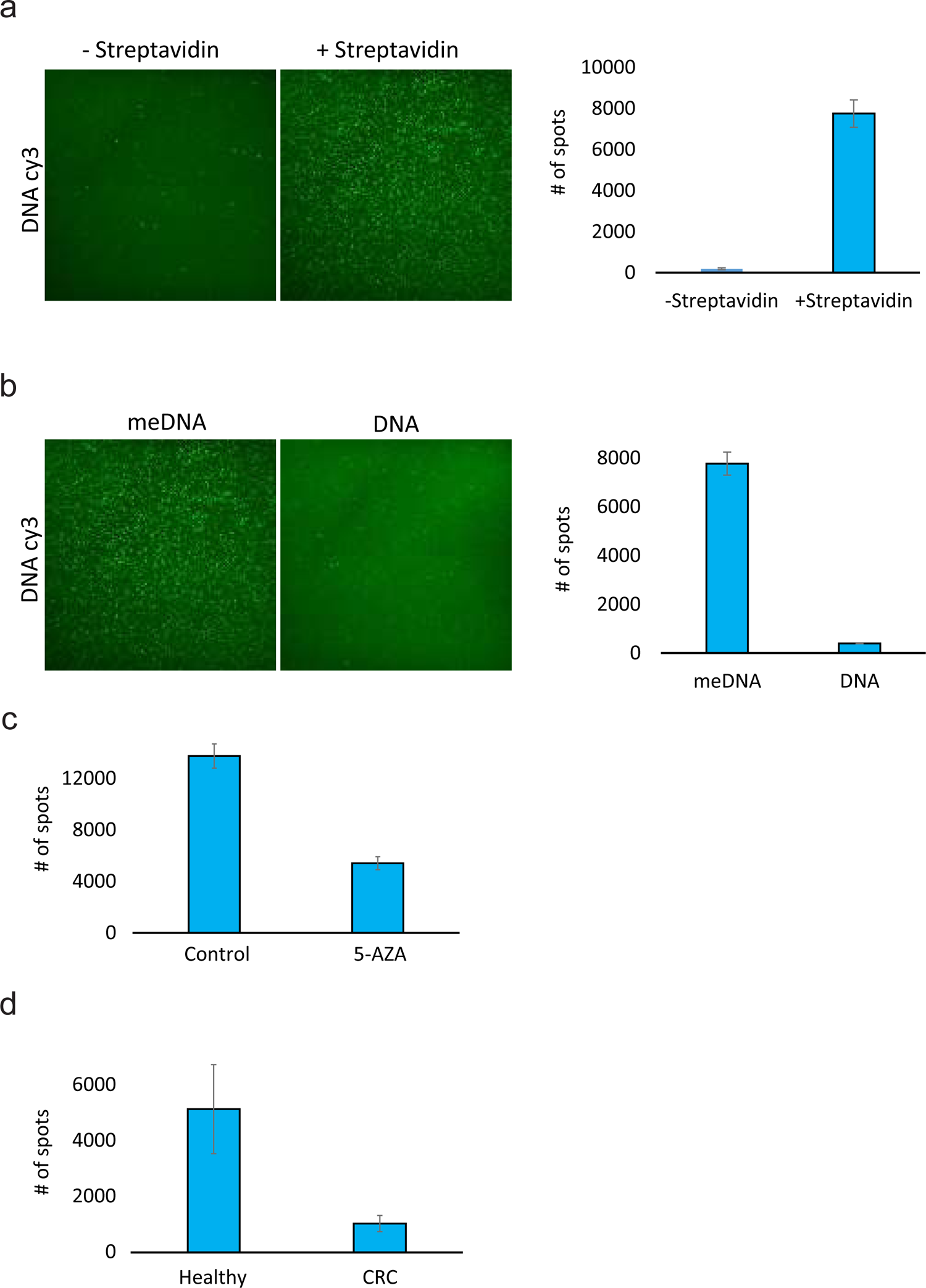
Single-molecule detection of DNA methylation levels. **(a)** Representative TIRF images (Left) and quantification (Right) indicating specific anchoring of Biotin-MBD-meDNA complexes to streptavidin-coated surfaces. Data is presented as the mean +/- s.d. of 50 FOV for each treatment. **(b)** Biotin-MBD specifically binds methylated DNA. Representative TIRF images (Left) and quantification (Right) of global DNA methylation levels following incubation of MBD2-biotin with Cy3-labeled (green) methylated (meDNA) or un-methylated synthetic DNA fragments. Data is presented as the mean +/- s.d. of 50 FOV for each treatment. **(c, d)** Quantification of global DNA methylation levels presented in Fig. 2i **(c)** and Fig. 2j **(d)**. Data is presented as the mean +/- s.d. of 50 FOV for each sample.

**Supplementary Fig.6.**
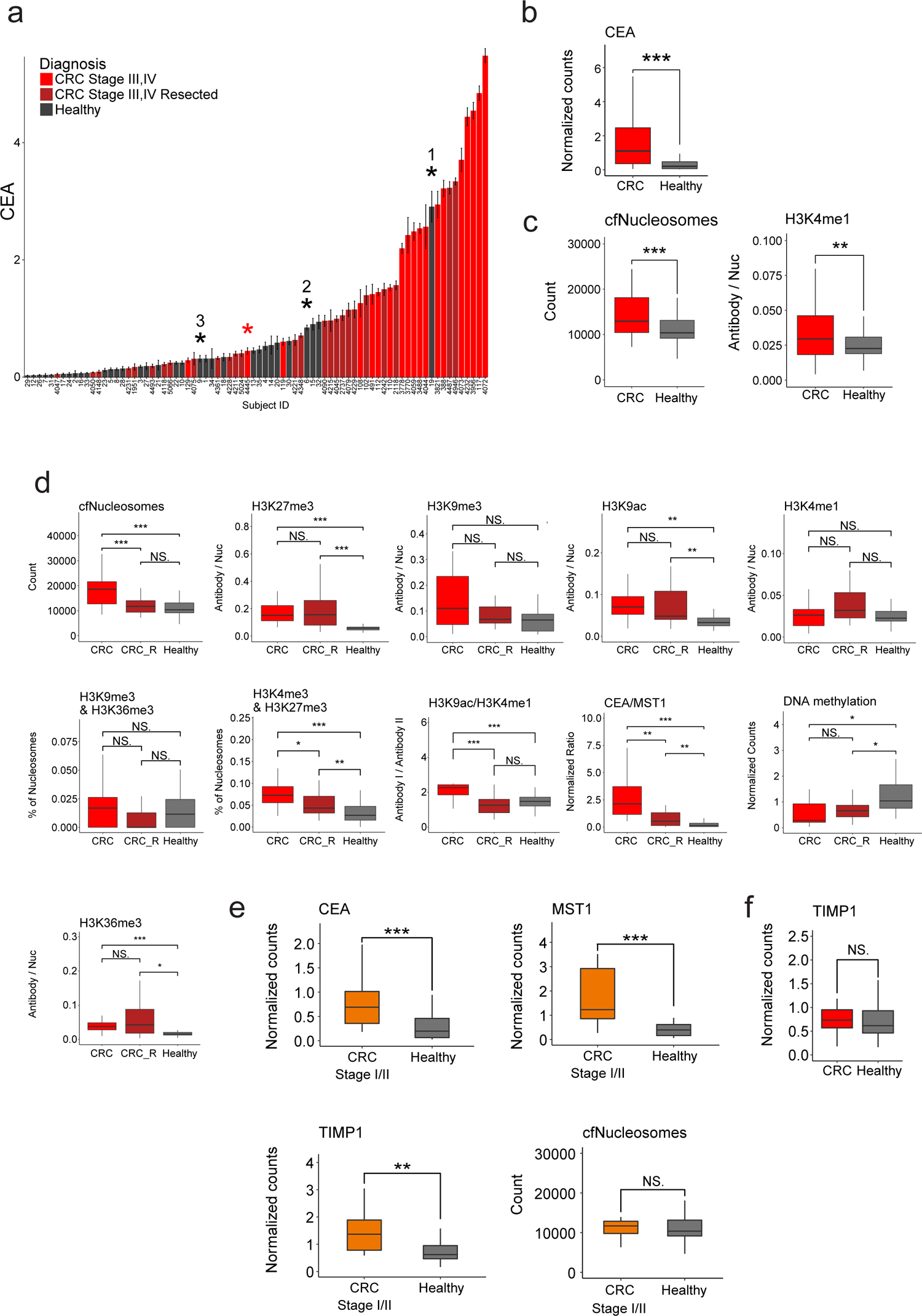
Analysis of histone PTMs, protein biomarkers and DNA methylation in the cohort of plasma samples from healthy and CRC subjects. **(a)** CEA normalized levels in the plasma of CRC patients and healthy individuals. Each bar represents a subject, data is presented as the mean +/- s.d. of 50 FOVs per sample. *1-3 correspond to healthy samples 19, 6, and 9, respectively (as in Fig. 3b). Sample 4445 (CRC, red) is denoted by *. **(b)** Box plot representation of the data in A (healthy = 33, CRC = 46(. Box plots limits: 25–75% quantiles, middle: median, upper (lower) whisker to the largest (smallest) value no further than 1.5× interquartile range from the hinge. P values were calculated by Welch’s t-test. *** P value < 0.001. **(c)** Global level of cfNucleosomes, and levels of H3K4me1-modified nucleosomes, significantly differ between healthy and CRC late stage samples (healthy = 33, CRC = 46(. P values were calculated by Welch’s t-test. * P value < 0.05 ** P value < 0.01. *** P value < 0.001. **(d)** Multiple comparison of all significant parameters between CRC, CRC resected (CRC_R) and healthy samples (CRC = 19, CRC_R=27, healthy = 33), corresponding to the data presented in main Fig. 3c-e. Of note, while all of these parameters differ between healthy versus the combined cohort of <u>all CRC</u> patients, this figure shows the differences between CRC patients with/without tumor resection. In some parameters, resected patients show higher similarity to healthy, and in other parameters, they are similar to CRC patients prior to tumor resection. See methods for P value calculation. * P value < 0.05 ** P value < 0.01. *** P value < 0.001. **(e)** Levels of CEA, MST1 and TIMP1 significantly differ between healthy and early stage CRC patients (healthy = 33, early CRC = 17). Total levels of cfNucleosomes do not show significant difference between the groups, likely due to low tumor burden at early stage patients. P values were calculated by Wilcoxon rank sum exact test. * P value < 0.05 ** P value < 0.01. *** P value < 0.001. **(f)** TIMP1 levels do not significantly differ in the cohort of healthy versus late-stage CRC patients.

**Supplementary Fig.7.**
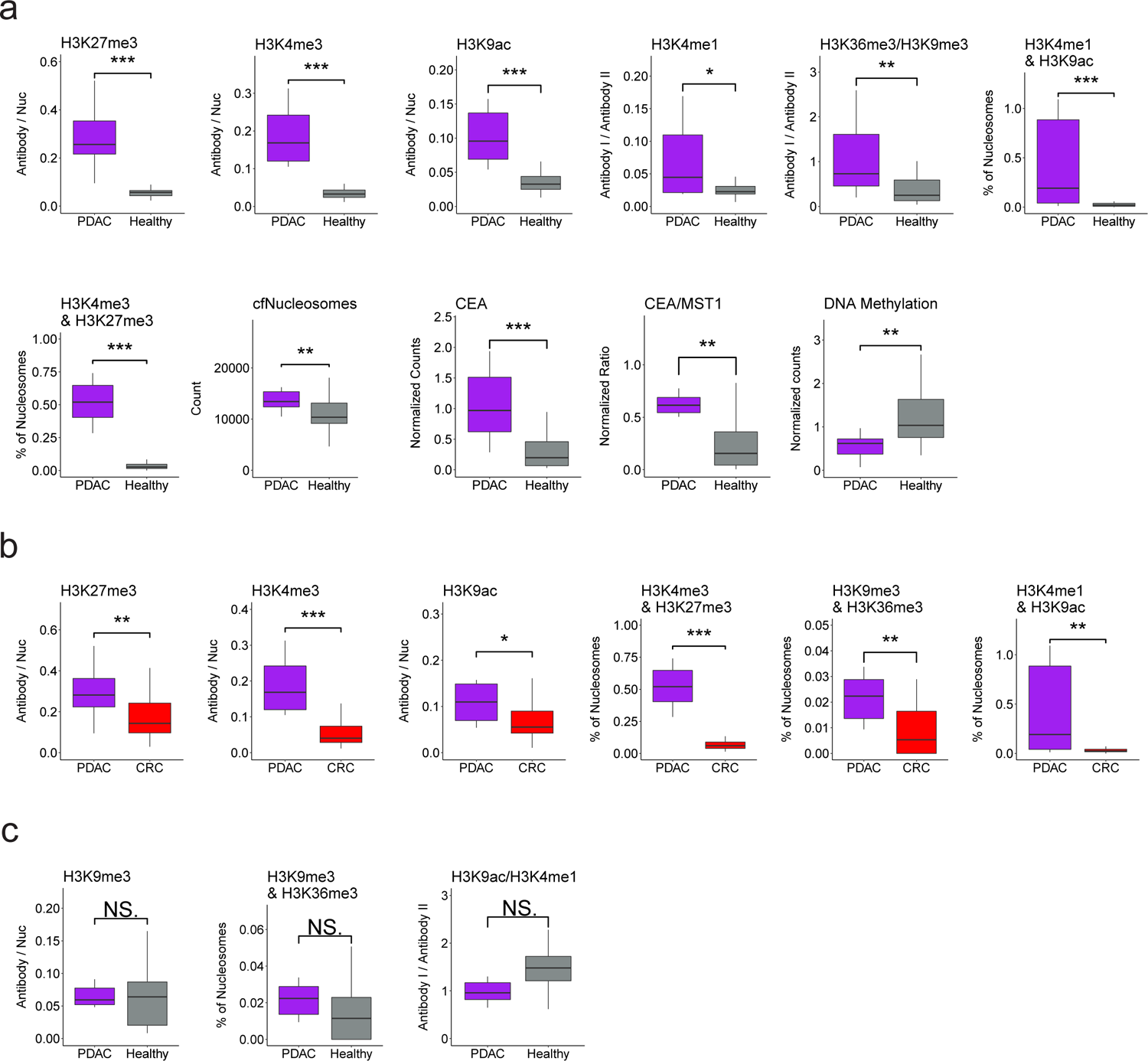
EPINUC reveals significant epigenetic and biomarkers alterations in the plasma of PDAC patients. **(a)** EPINUC measurements (as indicated on the graphs) that significantly differ between healthy and late stage PDAC samples (healthy = 33, PDAC = 10). P values were calculated by Wilcoxon rank sum exact test. * P value < 0.05 ** P value < 0.01. *** P value < 0.001. **(b)** EPINUC measurements (as indicated on the graphs) that significantly differ between PDAC and CRC late stage plasma samples (CRC = 46, PDAC = 10). P values were calculated by Wilcoxon rank sum exact test. * P value < 0.05 ** P value < 0.01. *** P value < 0.001. **(c)** Parameters that are specifically altered in late stage CRC compared to healthy (see Fig. 3d), but are not altered in patients diagnosed with PDAC. P values were calculated by Wilcoxon rank sum exact test. NS, Not significant.

**Supplementary Fig.8.**
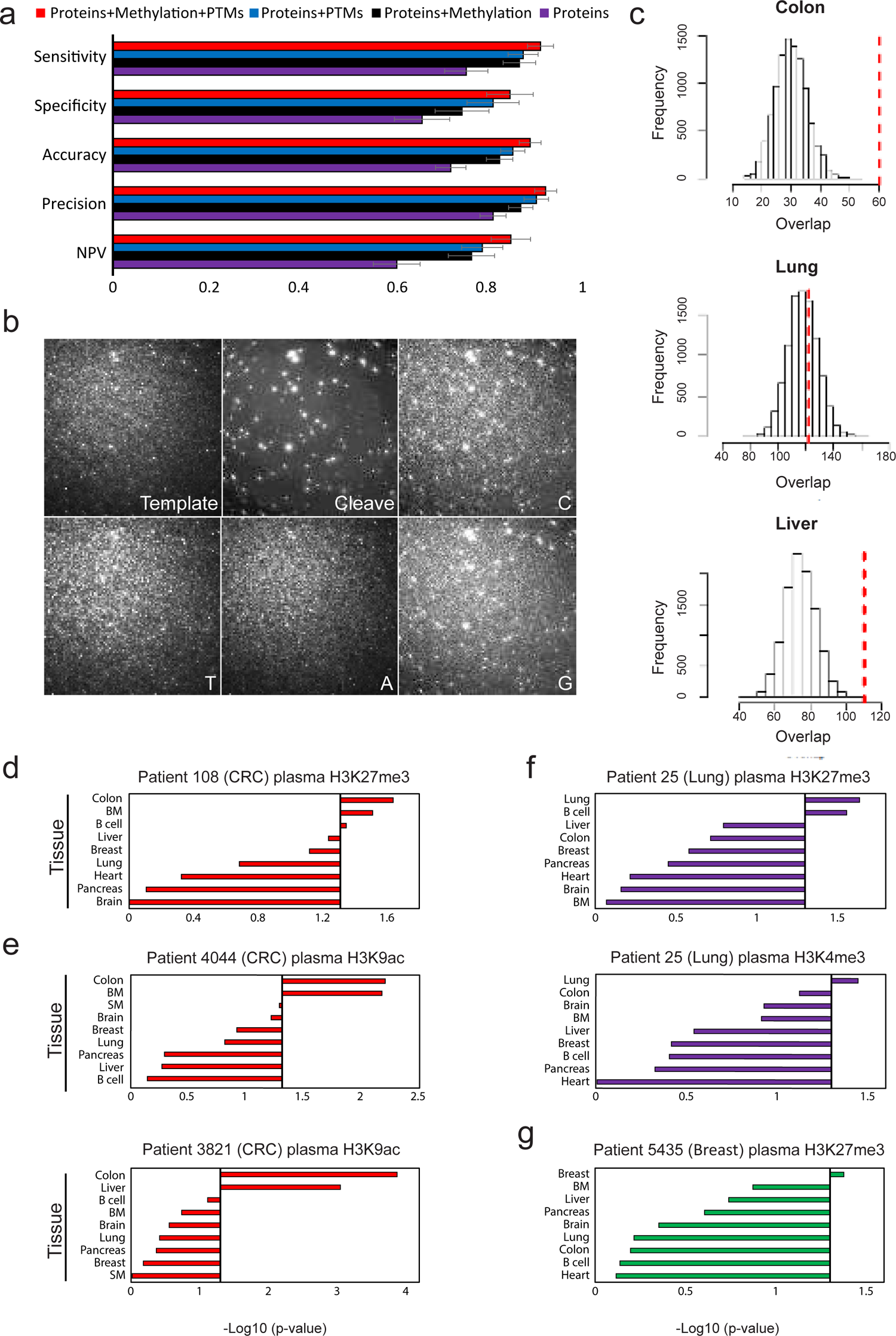
Prediction model performance and Tissue-of-origin analysis based on EPINUC-seq. **(a)** Logistic model performance increases with integration of additional layers of information. Corresponds to Fig. 4a. NPV, negative predictive value. **(b)** Representative TIRF images of antibodies’ binding (Template) followed by cleavage of fluorophore (Cleave) and sequencing cycles (C,T,A,G). High diameter spots correspond to FluoSpheres used for image alignment. **(c)** Histograms portray colon, lung and liver frequency distributions of random reads (equivalent to the number of H3K27me3 plasma CRC positive reads) that overlap with tissue H3K27me3 unique peaks (bootstrapping simulation, 10K iterations, Methods). Red line corresponds to the number of H3K27me3 unique tissue peaks that overlap with CRC (patient 4044) H3K27me3 positive plasma reads. **(d)** EPINUC-seq analysis of an additional CRC patient, similar to the analysis shown in Fig. 4d. **(e)** EPINUC-seq H3K9ac analysis of plasma from two CRC patients (stage IV). Tissues and primary cell lines ranked by overlap significance with single-molecule H3K9ac positive reads, indicating CRC as the tissue-of-origin. Heart H3K9ac peaks were available only for embryonic tissue (ENCODE), therefore were replaced with skeletal muscle tissue H3K9ac peaks. **(f)** EPINUC-seq analysis of plasma from stage IV lung cancer patient (patient 25). Tissues and primary cell lines ranked by overlap significance with single-molecule H3K27me3 (Left) or H3K4me3 (Right) positive reads, indicating lung as the tissue-of-origin. **(g)** EPINUC-seq analysis of plasma from stage IV Breast cancer patient (patient 5435). Tissues and primary cell lines ranked by overlap significance with single-molecule H3K27me3 positive reads, indicating breast as the tissue-of-origin. Black line corresponds to P value of 0.05. P values were determined by two tailed Z-test or Wilcoxon rank sum test.

**Supplementary Fig.9.**
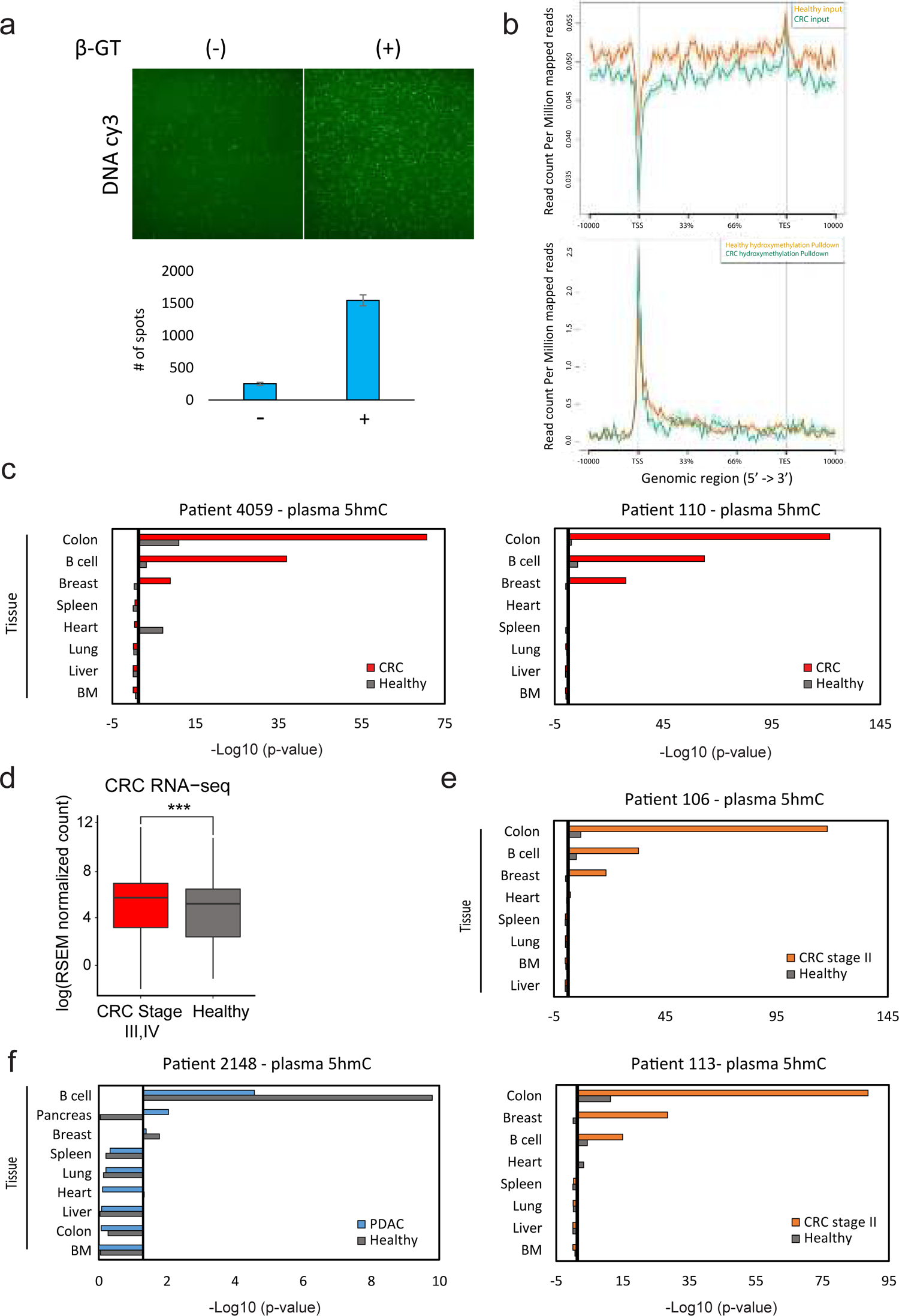
Tissue-of-origin analysis based on single-molecule 5hmC sequencing. **(a)** Representative TIRF images (Top) and quantification (Bottom) of fluorescently labeled nucleosomal DNA (Cy3, green) enriched for 5hmC, with (+) or without (-) the biotin conjugating enzyme beta-glucosyltransferase (β-GT). Data is presented as the mean +/- s.d. of 50 FOV for each treatment. **(b)** Metagene profiles of input (Top) and 5hmC enriched (Bottom) cfDNA sequenced from healthy (n=3, green) and CRC (n=3, Orange) samples, exhibiting 5hmC enrichment at gene bodies. **(c)** Overlap significance of tissues and primary cell lines unique H3K36me3 profiles with single-molecule 5hmC reads from healthy versus late stage CRC samples, similar to the analysis shown in Fig. 4g. **(d)** Boxplot depicts comparison of RNA expression in CRC primary tumor of each group of unique 5hmc gene signatures in healthy vs CRC (see Methods). RSEM, RNA-Seq by expectation-maximization. **(e-f)** Overlap significance of tissues and primary cell lines unique H3K36me3 profiles with single-molecule 5hmC reads from healthy versus early stage CRC **(e)** or healthy versus PDAC **(f)**, similar to the analysis shown in Fig. 4h. Each panel represents a different patient. Black line corresponds to P value of 0.05. P values were determined by Z-test.

**Supplementary Fig.10.**
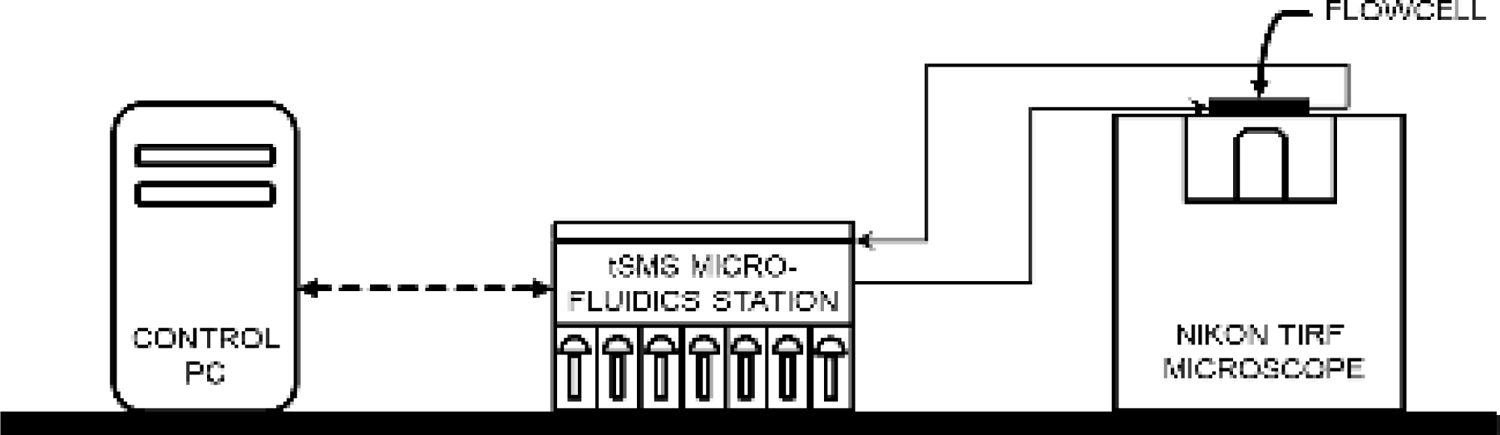
Single-molecule DNA sequencing technical setup. Schematic representation of key instrumentation used to perform single-molecule DNA sequencing. Straight and dashed lines represent microfluidic tubing and electric cables, respectively.

