## Supplementary table 2 for "Multiplexed Single-Molecule Epigenetic Analysis of Plasma-Isolated Nucleosomes for Cancer Diagnostics"

| **Antibody** |
|---|
| Anti-TIMP1Rabbit polyclonal (Biotin) |
| Anti-TIMP1 Rabbit polyclonal |
| MST1 Polyclonal Antibody |
| MST1 Monoclonal Antibody |
| CEA Mouse Monoclonal |
| CEA Mouse Monoclonal |
| p53 Mouse Monoclonal |
| Mutant p53 |
| p53 Rabbit Monoclonal (Alexa Fluor® 647 Conjugate) |
| Tri-Methyl-Histone H3 (Lys27) Rabbit mAb (AlexaFluor®488 Conjugate) |
| Acetyl-Histone H3 (Lys9) Rabbit mAb (Alexa Fluor®647 Conjugate) |
| Tri-Methyl-Histone H3 (Lys4) Rabbit mAb (In house Alexa Fluor®647 Conjugation) |
| Tri-Methyl-Histone H3 (Lys9) Rabbit mAb (Alexa Fluor® 488 Conjugate) |
| Mono-Methyl-Histone H3 (Lys4) XP®Rabbit mAb (Alexa Fluor® 488 Conjugate) |
| Tri-Methyl-Histon H3 (Lys36) XP(R)Rabbit mAb (Alexa Fluor® 647 Conjugate) |

| Vendor | Catalog |
| --- | --- |
| Abcam | ab77848 |
| Abcam | ab77847 |
| Thermo Fisher Scientific | PA587047 |
| Thermo Fisher Scientific | MA529824 |
| Thermo Fisher Scientific | MA117766 |
| Thermo Fisher Scientific | MIC0101 |
| Abcam | ab28 |
| Abcam | ab247264 |
| Abcam | ab227655 |
| CST | C36B11 |
| CST | C5B11 |
| CST | C42D8 |
| CST | D4W1U |
| CST | D1A9 |
| CST | D5A7 |
