## Supplementary table 4 for "Multiplexed Single-Molecule Epigenetic Analysis of Plasma-Isolated Nucleosomes for Cancer Diagnostics"

| Tissue/Primary cell | H3K27me3 Encode accession | |
|---|---|---|
| BM | ENCFF668KJG | ENCFF312GZL |
| B cell | ENCFF065LRV | ENCFF169OGM |
| Colon | ENCFF228JQE | ENCFF912DUE |
| Liver | ENCFF977LFS | ENCFF873VXC |
| Breast | ENCFF520QGK | ENCFF962YZN |
| Brain | ENCFF748JKZ | ENCFF878XDQ |
| Lung | ENCFF367LMT | ENCFF725NIC |
| Heart | ENCFF839XLE | ENCFF242JDO |
| Pancreas | ENCFF850CHF | ENCFF376GJG |

| Tissue/Primary cell | H3K4me3 Encode accession | |
|---|---|---|
| BM | ENCFF796GNT | ENCFF985RKZ |
| B cell | ENCFF763REX | ENCFF969DWP |
| Colon | ENCFF068YGR | ENCFF236IKZ |
| Liver | ENCFF076NWH | ENCFF178DYP |
| Breast | ENCFF562WCY | ENCFF590DGH |
| Brain | ENCFF410RJE | ENCFF724XKK |
| Lung | ENCFF254OZF | ENCFF526QUJ |
| Heart | ENCFF370ZUG | ENCFF437DEF |
| Pancreas | ENCFF095ENB | ENCFF642QJH |

| Tissue/Primary cell | H3K36me3 Encode accession | |
|---|---|---|
| BM | ENCFF802LQZ | ENCFF936ZZA |
| B cell | ENCFF157WLM | ENCFF769YLP |
| Colon | ENCFF732NMQ | ENCFF893YTR |
| Liver | ENCFF220MSF | ENCFF342YOA |
| Breast | ENCFF540RWM | ENCFF648VTD |
| Spleen | ENCFF380PDF | ENCFF472BWO |
| Lung | ENCFF597AKE | ENCFF829QFZ |
| Heart | ENCFF224FAC | ENCFF912SFR |
| Pancreas | ENCFF267LPH | ENCFF283YWD |

| Tissue/Primary cell | H3K9ac Encode accession | |
|---|---|---|
| BM | ENCFF302LTS | NA |
| B cell | ENCFF181FLS | NA |
| Colon | ENCFF023PFF | ENCFF486TGX |
| Liver | ENCFF344IUW | ENCFF766JJU |
| Breast | ENCFF152HYX | ENCFF294GAH |
| Brain | ENCFF507ZNV | ENCFF769GHM |
| Lung | ENCFF635CCP | ENCFF958SIQ |
| Skeletal Muscle | ENCFF307JWI | ENCFF310QAQ |

| | | |
|---|---|---|
| Plasma Healthy | Healthy 1-3 | Data accquired from Sadeh et al. 2021 |
| Plasma CRC | CRC patients 1-3 | Data accquired from Sadeh et al. 2021 |
